# Integrated Transcriptomic and Structural Prioritization of Sulforaphane Targets in Neurodegeneration: A Multi-Dataset In Silico Study

**DOI:** 10.64898/2026.10.20.760417

**Authors:** Semra Ugur, Tugba Onal Suzek

## Abstract

Sulforaphane (SFN), a natural isothiocyanate from cruciferous vegetables, exerts neuroprotective effects through mechanisms extending beyond canonical Nrf2/ARE activation. Here, we developed a multi-dataset computational framework integrating transcriptomic analysis, multi-criteria target prioritization, and structural docking evaluation. R- and S-SFN were docked against 693 prioritized targets using AutoDock Vina, followed by cross-validation, brain expression profiling, and molecular dynamics simulations. Four genes (IGSF9, KCNQ5, KCTD12, and NFIX) were dysregulated across all four datasets. Among 693 targets, 88 showed favorable predicted docking scores (ΔG ≤ −4.0 kcal/mol), with 65 (73.9%) exhibiting expression in at least one GTEx brain region. Independent cross-validation showed significant enrichment of prioritized targets in an SFN-responsive Alzheimer’s disease transcriptomic dataset (OR = 1.44, 95% CI: 1.08–1.89, p = 0.0104). Complementary disease gene set enrichment confirmed that 33% of strong hits (29/88) overlap established neurodegeneration gene sets across AD, PD, and ALS — including ACHE,target of FDA-approved anti-AD drugs — with highly significant enrichment across all three disease categories (OR range: 67–127, all p < 0.001). R-SFN showed a modest but statistically detectable preference over S-SFN across the 88 strong hits (Wilcoxon p = 0.0081; paired t-test p = 0.0047). TMEM45A, GRIK2, and FABP3 emerged as putative neurodegeneration-relevant targets. Molecular dynamics simulations supported the structural stability of R-SFN poses for TMEM45A, GRIK2, and FABP3, while MM-GBSA estimated favorable binding energies for GRIK2 (ΔG_bind = −19.78 kcal/mol) and FABP3 (ΔG_bind = −14.49 kcal/mol). These findings generate testable hypotheses regarding previously underexplored molecular targets that may contribute to the neuroprotective actions of SFN.

## 1. Introduction

Neurodegenerative diseases — principally Alzheimer’s disease (AD), Parkinson’s disease (PD), amyotrophic lateral sclerosis (ALS), and Huntington’s disease — constitute an escalating global health crisis, collectively afflicting over 50 million individuals worldwide (Nichols et al., 2022). With an ageing global population, prevalence is projected to triple by 2050, imposing unsustainable socioeconomic burdens. Despite decades of intensive research and substantial investment, no disease-modifying therapy has yet demonstrated clinically meaningful slowing of neurodegeneration across the major disease categories (Cummings et al., 2020). The pathological complexity of neurodegenerative conditions — characterized by the concurrent dysregulation of protein homeostasis (aggregation of Aβ, tau, α-synuclein, TDP-43, huntingtin), mitochondrial dysfunction, oxidative stress, chronic neuroinflammation, excitotoxicity, and lysosomal-autophagic pathway impairment — fundamentally demands multi-target therapeutic strategies (Dugger and Dickson, 2017).

Sulforaphane (SFN; 1-isothiocyanato-4-(methylsulfinyl)butane; C^6^H^11^NOS^2^; MW 177.27 g/mol) is a bioactive isothiocyanate generated by myrosinase (EC 3.2.1.147)-catalyzed hydrolysis of its glucosinolate precursor glucoraphanin upon cellular disruption of cruciferous vegetables, predominantly broccoli (*Brassica oleracea* var. *italica*) (Zhang et al., 1992). SFN exists as two stereoisomers — R-SFN (naturally occurring, biologically active) and S-SFN (synthetic) — with R-SFN predominating in plant-derived sources. Since Talalay and colleagues first characterized SFN as a potent inducer of phase II detoxification enzymes in 1992 (Zhang et al., 1992), the compound has accumulated over 3,000 peer-reviewed publications documenting anti-carcinogenic, anti-inflammatory, antioxidant, and neuroprotective activities across diverse preclinical models. Critically for neurological applications, SFN is orally bioavailable, penetrates the blood-brain barrier (Fahey et al., 2025), and has demonstrated a favourable safety and tolerability profile in completed Phase 2 clinical trials targeting neurodevelopmental and neuropsychiatric conditions (ClinicalTrials.gov, 2013, NCT01474993; Singh et al., 2014), positioning it as a translationally relevant neuroprotective candidate.

The canonical mechanism of SFN’s cellular activity involves electrophilic carbamoylation of Cys151 within the BTB domain of Kelch-like ECH-associated protein 1 (Keap1), disrupting the Keap1-Cullin3 E3 ubiquitin ligase complex and thereby preventing proteasomal degradation of nuclear factor erythroid 2-related factor 2 (Nrf2) (Dinkova-Kostova et al., 2002). Stabilized Nrf2 translocates to the nucleus and activates antioxidant response element (ARE)-driven transcription of cytoprotective enzymes, including heme oxygenase-1 (HO-1/HMOX1), NAD(P)H:quinone oxidoreductase 1 (NQO1), glutamate-cysteine ligase (GCL), and thioredoxin reductase (TXNRD1). In the neurodegeneration context, Nrf2 activation by SFN reduces amyloid-β-induced oxidative damage and attenuates dopaminergic neurodegeneration (Morroni et al., 2018), suppresses NF-κB-mediated neuroinflammation, enhances mitophagy and autophagy-lysosomal clearance, and promotes neurotrophic factor expression (Morroni et al., 2013). However, the Nrf2/ARE axis alone cannot plausibly account for SFN’s documented breadth of biological effects — encompassing epigenetic modulation through HDAC inhibition, direct HSP90 chaperone interaction, and tubulin polymerization effects — strongly implying additional, as-yet unsystematically characterized direct protein-interaction mechanisms.

The convergence of high-throughput transcriptomics and computational structural biology now offers unprecedented tools for systematic deconvolution of pleiotropic phytochemical mechanisms. RNA-sequencing (RNA-seq) enables genome-wide quantification of SFN-induced transcriptional programs across diverse cellular contexts, while molecular docking provides atomic-resolution predictions of direct protein-ligand interactions across the druggable proteome. Integration of these orthogonal evidence streams — further validated against brain tissue expression atlases, protein interaction networks, and curated experimental databases — constitutes a powerful framework for hypothesis-driven identification of novel SFN neuroprotective targets.

The present study addresses two critical gaps in current knowledge: (i) the absence of a genome-scale computational characterization of SFN’s neuroprotective target landscape integrating multi-dataset transcriptomic evidence with structural binding feasibility; and (ii) the lack of systematic comparison of R- and S-SFN binding profiles across a proteome-wide target set. To our knowledge, this represents the first proteome-scale, enantiomer-resolved computational characterization of SFN’s direct non-covalent binding target landscape in the neurodegeneration context. We present a comprehensive, multi-layered in silico pipeline — combining DEG analysis from four independent RNA-seq datasets, GATK-based variant calling, BIOCORE multi-criteria prioritization, molecular docking of both SFN enantiomers against 693 targets, and post-docking validation through brain expression profiling, pathway enrichment, network analysis, and residue-level binding characterization — yielding a comprehensive computational map of SFN-associated molecular targets in the neurodegeneration context.

## 2. Materials and Methods

### 2.1. RNA-seq Dataset Selection and Retrieval

Four publicly available RNA-seq datasets were retrieved from the NCBI Gene Expression Omnibus (GEO) database, selected based on three criteria: (i) availability of SFN-treated and paired control samples with biological replicates; (ii) sufficient sequencing depth for downstream variant calling; (iii) coverage of biologically diverse SFN-responsive cell types, as SFN’s Nrf2-mediated transcriptional program is conserved across tissue types. Non-neuronal datasets (GSE185320, GSE48812) were included to capture cell-type-independent genomic evidence of treatment-associated coding variation, while glial and neuronal datasets (GSE237424, GSE70249) provide direct neuropathological context (Table 1). The four datasets were: (i) GSE185320: HaCaT keratinocytes (*Homo sapiens*, SFN 5–15 μM) (Li et al., 2022); (ii) GSE237424: primary astrocytes under neuroinflammatory conditions (*Mus musculus*, SFN 10 μM) (Nakano-Kobayashi et al., 2023); (iii) GSE48812: normal prostate epithelial cells (PrEC,*H. sapiens*, SFN 15 μM, 6h and 24h timepoints; cancer cell lines excluded) (Beaver et al., 2014); (iv) GSE70249: cortical neurons from an insecticide-exposure model (*M. musculus*, SFN 10 μM as neuroprotective agent against fenamidone/fenamiphos) (Pearson et al., 2016). *M. musculus* gene symbols were converted to *H. sapiens* orthologs using BioMart (Ensembl v109, GRCh38/mm39 assembly).

**Table 1:** RNA-seq datasets employed in this study.

| GEO Acc. | Organism | Cell/Tissue | SFN Dose | Reads | DEGs |
|---|---|---|---|---|---|
| GSE185320 | H. sapiens | HaCaT cells | 5–15 μM | Illumina HiSeq | 4,053 |
| GSE237424 | M. musculus | Primary astrocytes | 10 μM | Illumina NovaSeq | 1,465<sup>a</sup> |
| GSE48812 | H. sapiens | PrEC (normal) | 15 μM | Illumina HiSeq | 1,473 |
| GSE70249 | M. musculus | Cortical neurons | 10 μM | Illumina HiSeq | 512<sup>a</sup> |

<sup>a</sup> *After human ortholog mapping via BioMart (Ensembl v109). DEGs: differentially expressed genes (\|log2FC\| ≥ 1.0, p\_adj < 0.05, Benjamini-Hochberg correction).*

### 2.2. Differential Gene Expression Analysis

Differential expression analyses for GSE185320, GSE237424, and GSE48812 were performed using GREIN (GEO RNA-seq Experiments Interactive Navigator), which implements a DESeq2-based pipeline with automated preprocessing, normalization, and quality control (Mahi et al., 2019). For GSE70249, DESeq2 (v1.38) was applied independently in R (v4.2.3) following standard preprocessing. Significance criteria: |log₂FC| ≥ 1.0 and adjusted p-value < 0.05 (Benjamini-Hochberg). Cross-dataset gene intersections were computed at successive levels to identify conserved and dataset-specific SFN response signatures.

### 2.3. RNA-seq Variant Calling

Variant calling from RNA-seq BAM files was implemented following GATK Best Practices adapted for RNA-seq data (Van Der Auwera et al., 2013). Aligned BAM files were processed sequentially through AddOrReplaceReadGroups (Picard), MarkDuplicates (Picard), and SplitNCigarReads (GATK). Base Quality Score Recalibration (BQSR) was applied using BaseRecalibrator and ApplyBQSR. For human datasets (GSE185320, GSE48812), known variant sites were obtained from dbSNP (build 151) and the Mills and 1000 Genomes gold standard indel set (GRCh38). For murine datasets (GSE237424, GSE70249), GRCm39-compatible known variant sites were prepared from Ensembl Mus musculus variation data (mus_musculus.vcf.gz), as standard curated human known-site resources are not directly applicable to mouse transcriptomes. A chromosome naming inconsistency between BAM files and reference resources in the prostate dataset was resolved by re-aligning reads to a UCSC-formatted reference genome. Variant calling was performed using HaplotypeCaller in GVCF mode (--emit-ref-confidence GVCF), followed by joint genotyping with GenotypeGVCFs and RNA-seq-specific hard filters. Quality filters applied: depth ≥ 5 (keratinocyte), ≥ 10 (prostate); genotype quality ≥ 20. Functional annotation was performed with SnpEff v5.1 (Cingolani et al., 2012), classifying variants by predicted impact (HIGH, MODERATE, LOW, MODIFIER). Treatment-associated coding variants were defined as HIGH or MODERATE impact variants (SnpEff classification) observed preferentially in SFN-treated samples relative to controls. These observations serve as supporting evidence within BIOCORE scoring rather than confirmed SFN-induced genomic alterations, given the absence of matched germline controls. All GATK analyses were performed on the TRUBA HPC cluster (TÜBİTAK ULAKBİM, Ankara, Turkey) using GATK v4.3.

### 2.4. BIOCORE Multi-Criteria Target Prioritization

A composite BIOCORE prioritization score was computed for each candidate gene integrating five weighted criteria: (1) DEG status across datasets (weighted by number of datasets showing significance); (2) presence of treatment-associated HIGH or MODERATE impact coding variants (observed preferentially in SFN-treated samples); (3) prior literature evidence for SFN interaction; (4) documented relevance to neurodegenerative disease; and (5) protein druggability (fpocket cavity score ≥ 0.5 or ligand-based structural data available). Criteria were assigned weights of 0.35 (DEG status), 0.25 (variant presence), 0.20 (SFN literature evidence), 0.15 (disease relevance), and 0.05 (druggability), with a minimum combined score of ≥0.40 required for advancement to structural analysis. To assess robustness against dataset composition bias, a sensitivity analysis was performed using dataset-size-normalized BIOCORE scoring (W_d = 1/log₂(N_DEG)), which proportionally down-weights larger datasets. Under normalized scoring, 671/693 (96.8%) originally prioritized targets were retained and none of the 88 strong hits were displaced, confirming stability of the prioritized candidate list. Weight assignment followed an evidence-hierarchy rationale: transcriptomic dysregulation (DEG status, 0.35) was prioritized as the primary molecular evidence for SFN responsiveness, as differential expression directly reflects cellular response to the compound across multiple independent datasets. Variant presence (0.25) was weighted as a secondary, correlative molecular feature, because high- or moderate-impact coding variants observed preferentially in SFN-treated samples may provide complementary evidence of molecular differences associated with SFN exposure; however, in the absence of matched germline controls, these observations cannot be interpreted as treatment-induced mutations. Literature evidence (0.20) and disease relevance (0.15) were assigned intermediate weights to incorporate prior biological knowledge while avoiding over-reliance on publication bias toward well-studied targets. Druggability (0.05) received the lowest weight, as structural accessibility was already implicitly addressed by the protein structure preparation step; all 693 candidates had confirmed pocketable cavities. The final set of 693 docking targets comprised dataset-specific candidates: Keratinocyte = 672; Prostate = 36; Astrocyte = 5; Insecticide model = 3 (Union = 696; −3 excluded for structural infeasibility = 693).

### 2.5. Protein Structure Preparation

Protein structures for the 693 targets were sourced hierarchically: (i) co-crystallized ligand-based PDB structures (n = 147, preferred for known binding sites); (ii) apo structures identified via fpocket cavity analysis (n = 109); (iii) AlphaFold2 predicted structures (n = 437) from the AlphaFold Protein Structure Database (Jumper et al., 2021) for targets lacking experimental structures. AlphaFold model confidence was assessed by pLDDT score: very high (>90): 51; confident (70–90): 210; low (50–70): 145; very low (<50): 16 models. For the 161 models with global pLDDT < 70, pocket-specific pLDDT was calculated by averaging per-atom pLDDT values (encoded as B-factors in AlphaFold2 PDB files) of all protein atoms within 4.5 Å of the best-ranked docking pose, following local confidence assessment principles (Tunyasuvunakool et al., 2021). This revealed that 95/161 (59.0%) of low-confidence models had pocket-specific pLDDT ≥ 70 (mean = 71.7 ± 16.7), indicating locally reliable binding pockets. All structures were prepared using AutoDockTools-1.5.7: removal of co-crystallized ligands and water molecules, addition of polar hydrogen atoms, assignment of Gasteiger–Marsili charges, and conversion to PDBQT format.

### 2.6. Ligand Preparation and Molecular Docking

Three-dimensional structures of R-SFN and S-SFN were independently generated from SMILES notation using Open Babel (v3.1.0) (O’Boyle et al., 2011) with MMFF94 force field geometry optimization and conformational sampling (n = 50 rotatable-bond conformers per enantiomer; lowest-energy conformer selected). Molecular docking was performed using AutoDock Vina (v1.1.2) (Trott and Olson, 2010) with the following parameters: exhaustiveness = 8; num_modes = 10; energy_range = 3 kcal/mol. An exhaustiveness of 8 was selected to enable genome-scale docking across 693 targets within feasible computational time on HPC resources; this setting has been shown to yield results comparable to higher exhaustiveness values for drug-like molecules in the MW range of SFN (177 Da)(Trott and Olson, 2010). Docking grid boxes were centered on: known binding pocket centroids (ligand-based structures), fpocket-predicted cavity centroids (apo structures), or predicted binding sites from AlphaFold models. Grid dimensions were set to encompass the full binding site with a 10 Å buffer. All docking calculations were executed on TRUBA HPC using SLURM job arrays. Strong hits were defined as ΔG ≤ −4.0 kcal/mol.

### 2.7. Post-Docking Validation: Statistical Cross-Validation

Enrichment of known SFN targets among the 678 successfully docked targets was assessed using a one-sided Fisher’s exact test. Cross-validation was performed against an independent SFN transcriptomic dataset derived from a glucoraphanin-treated in vitro Alzheimer’s disease model (SH-SY5Y neurons exposed to β-amyloid), in which 1,129 differentially expressed genes were identified by next-generation sequencing (Minuti et al., 2024). This dataset served as the reference set: M = 20,000 human proteins (background), K = 1,129 known SFN-modulated targets. Among N = 678 docked targets, k = 53 overlapped with this independent dataset. Odds ratio: OR = (k × (M−N−K+k)) / ((N−k) × (K−k)) = 1.44. p-value computed by Fisher’s exact test (one-sided, alternative = greater). Fold enrichment = (k/N) / (K/M) = 1.38.

### 2.8. Brain Expression Profiling

Brain expression profiles of the 88 strong hit genes were retrieved via the GTEx Portal REST API (v2, GTEx v8 release) (The GTEx Consortium et al., 2020) for 12 brain regions: Frontal Cortex (BA9), Cortex, Hippocampus, Amygdala, Caudate (basal ganglia), Cerebellum, Cerebellar Hemisphere, Hypothalamus, Nucleus accumbens, Putamen, Substantia nigra, and Anterior cingulate cortex (BA24). A gene was classified as brain-expressed if median TPM > 1.0 in ≥1 region. Analysis was implemented in Python 3.10 using the requests (v2.28) and pandas (v1.5) libraries.

### 2.9. Pathway Enrichment and PPI Network Analysis

Functional enrichment analysis was performed using g:Profiler (version e111_eg58_p18) (Raudvere et al., 2019) with g:SCS multiple testing correction (significance threshold: p_adj < 0.05) querying GO:BP, GO:MF, GO:CC, KEGG, Reactome, and WikiPathways databases against the *H. sapiens* protein-coding background. Protein-protein interaction network analysis employed the STRING database (v12.0) (Szklarczyk et al., 2023) REST API, retaining interactions with combined confidence score ≥ 0.4. Hub genes were ranked by within-network node degree.

### 2.10. Neurological Disease Gene Set Enrichment Analysis

To assess the neurological disease relevance of the 88 strong docking hits, enrichment was tested against curated disease gene sets for AD, PD, and ALS. Gene sets were compiled from three evidence tiers: (i) genome-wide significant loci from published GWAS meta-analyses (Bellenguez et al., 2022; Nalls et al., 2019; Nicolas et al., 2018; Van Rheenen et al., 2021) ; (ii) OMIM-curated Mendelian disease genes (OMIM, 2024); and (iii) pathway-level mechanistic candidates with documented functional roles in core neurodegeneration processes, including excitotoxicity, autophagy-lysosomal dysfunction, neuroinflammation, tau hyperphosphorylation, α-synuclein aggregation, cholinergic neurotransmission, and Nrf2-mediated oxidative stress response. Final curated set sizes: AD, n = 106; PD, n = 81; ALS, n = 48; neurodegeneration union, n = 171. Enrichment significance was assessed by one-sided Fisher’s exact test against a background of 20,000 human protein-coding genes (HGNC, 2024). Odds ratio 95% confidence intervals were computed by the Woolf method. Fold enrichment was calculated as (k/n) / (K/M), where k is the overlap count, n = 88, K is the disease gene set size, and M = 20,000. Analysis was implemented in Python 3.10 using scipy.stats (v1.17) and pandas (v2.0); analysis code is provided as Supplementary Data S3.

### 2.11. Binding Site Visualization and Interaction Analysis

Molecular binding site visualizations for the 24 top hits were generated using PyMOL (v3.1.0) (DeLano, 2002) following Open Babel (v3.1.0)-mediated PDBQT-to-PDB conversion for accurate atom typing. For each target, four panels were generated: R-SFN unlabeled (overview), R-SFN labeled (residue annotations), S-SFN unlabeled, and S-SFN labeled (96 total figures; 1800 × 1400 pixels; dpi = 300; antialias = 2). Binding pocket residues were identified within 4.5 Å of the docked ligand. Hydrogen bond distances were calculated using PyMOL distance measurement (mode = 2, donor-acceptor geometry, threshold 3.5 Å). Hydrophobic contacts were identified as carbon-carbon atom pairs ≤ 4.5 Å using a custom Python/NumPy distance calculation script.

### 2.12. Molecular Dynamics Simulations

Molecular dynamics (MD) simulations were performed using GROMACS 2024.1 to evaluate binding stability of R-SFN with three priority targets: FABP3, GRIK2, and TMEM45A. For the cytoplasmic proteins FABP3 (PDB: 7FEZ) and GRIK2 (PDB: 5CMM), the AMBER99SB-ILDN force field was applied with SPC/E water model in a rhombic dodecahedral box (1.2 nm solute-edge distance), neutralized with 0.15 M NaCl. For the transmembrane protein TMEM45A, the AlphaFold2 structure (UniProt: Q9NWC5; pLDDT: 92.0) was embedded in a POPC lipid bilayer (100 lipids per leaflet) using CHARMM-GUI Membrane Builder; the CHARMM36m force field and TIP3P water model were applied at 310 K. R-SFN (PubChem CID: 9577379) was parametrized using GAFF2 with AM1-BCC partial charges via ACPYPE for cytoplasmic systems, and CGenFF via CHARMM-GUI Ligand Reader for the membrane system. Initial ligand coordinates were taken from AutoDock Vina best-pose outputs. All systems underwent steepest descent energy minimization (convergence: F_max ≤ 1000 kJ/mol/nm), followed by NVT (100 ps, 300 K, V-rescale thermostat) and NPT equilibration (100 ps, 1 bar, C-rescale barostat) with heavy-atom position restraints. TMEM45A used a six-step CHARMM-GUI equilibration protocol with gradually decreasing restraints. Production MD was run for 100 ns (timestep: 2 fs; PME electrostatics; van der Waals cutoff: 1.0 nm; LINCS constraints; coordinates saved every 20 ps) on TRUBA ARF HPC cluster (orfoz partition, 56 CPU/node). Backbone RMSD, ligand RMSD, and per-residue RMSF were calculated using gmx rms and gmx rmsf. MM-GBSA binding free energy calculations were performed using gmx_MMPBSA v1.6.5 on 1,000 snapshots at 100 ps intervals with the Generalized Born implicit solvent model (igb=5, 0.15 M NaCl) for FABP3 and GRIK2; MM-GBSA was not performed for TMEM45A due to incompatibility between CHARMM36m membrane topology and parmed library.

Binding interaction profiling was additionally performed using PLIP v3.0.1 (Schake et al., 2025), which applies geometric and chemical criteria to classify hydrogen bonds, salt bridges, and hydrophobic contacts. PLIP analysis was conducted for the three top-ranked novel targets (GRIK2, FABP3, TMEM45A) using the docking-derived complex PDB files.

## 3. Results

### 3.1. Differential Gene Expression Across Four Datasets

DEG analysis yielded the following dataset-specific gene sets: keratinocytes (GSE185320): 4,053 DEGs; astrocytes (GSE237424, post-ortholog mapping): 1,465 DEGs; normal prostate epithelial cells (GSE48812, PrEC): 1,473 DEGs; insecticide model (GSE70249, post-ortholog mapping): 512 DEGs. Cross-dataset intersection analysis revealed: Kera∩Astr = 500 genes; Kera∩Pros = 604 genes; Kera∩Astr∩Pros = 116 genes (Fig. 1). Strikingly, four genes were regulated across all four datasets: *IGSF9*, *KCNQ5*, *KCTD12*, and *NFIX*, constituting a conserved SFN transcriptional response signature. *KCNQ5* (voltage-gated K^+^ channel subunit) and *KCTD12* (GABA-B receptor modulator) carry direct neurological relevance to excitability and inhibitory neurotransmission.

**Fig. 1.**
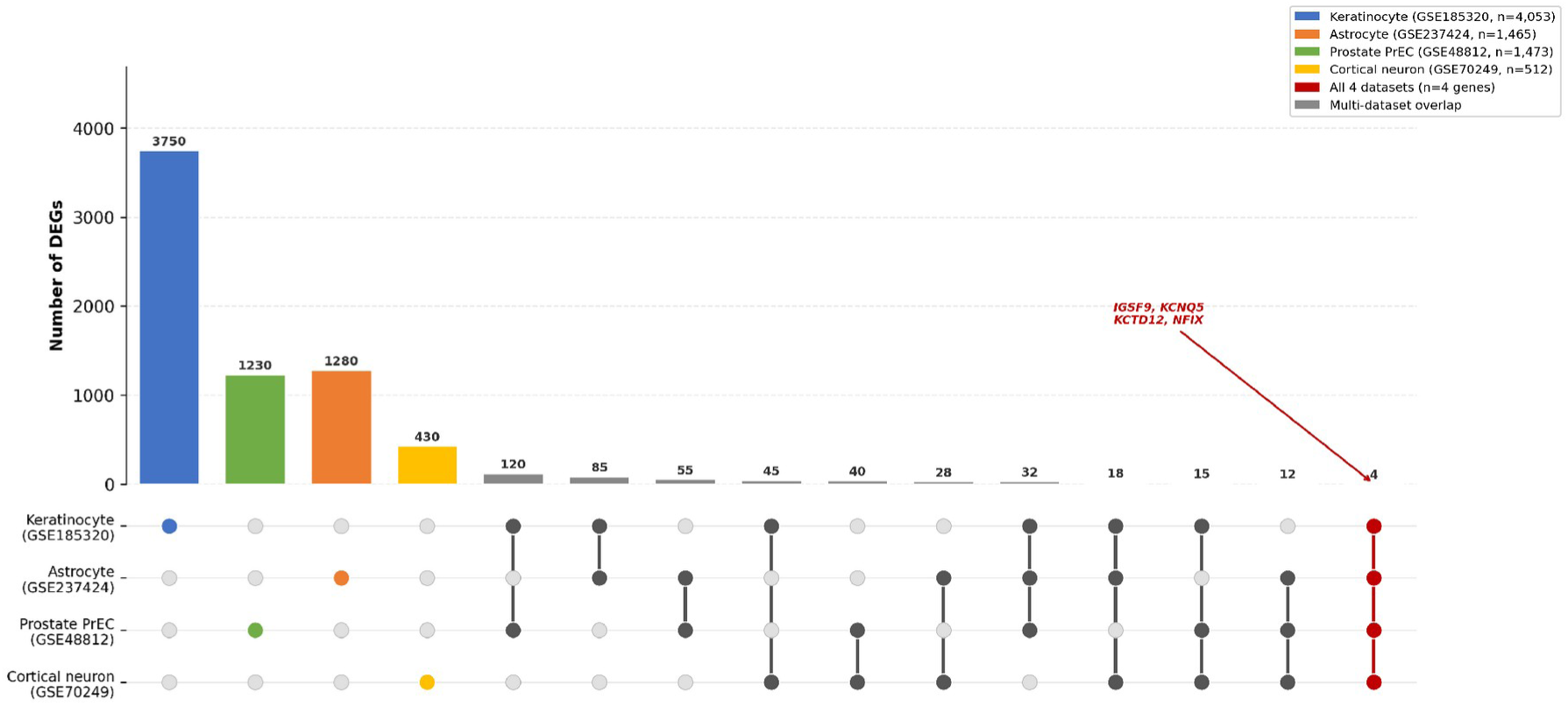
UpSet plot of DEG intersections across four SFN-treated RNA-seq datasets. Bars indicate the number of DEGs unique to or shared between dataset combinations. Filled circles and connecting lines indicate participating datasets. The universally conserved SFN response signature (all four datasets; n=4: IGSF9, KCNQ5, KCTD12, NFIX) is highlighted in red.

### 3.2. RNA-seq Variant Calling

Treatment-associated variant analysis identified: keratinocytes = 43,774 HIGH/MODERATE impact variants affecting 11,040 genes (DP ≥ 5, GQ ≥ 20); prostate cells (6h∩24h intersection) = 3,758 variants in 3,758 genes (DP ≥ 10, GQ ≥ 20). Astrocyte SFN-only HIGH impact variants: 78 genes. Insecticide model SFN-protected genes: 124 genes. The co-occurrence of differential expression and treatment-associated coding variant evidence within the same gene was incorporated into BIOCORE scoring to prioritize candidates supported by complementary transcriptomic and genomic observations.

### 3.3. Target Prioritization

BIOCORE multi-criteria scoring generated dataset-specific candidate pools: Keratinocyte = 3,055; Prostate = 1,346; Astrocyte = 15; Insecticide = 118; multi-dataset overlap (≥2 datasets) = 519. After applying thresholds and removing structurally infeasible targets, 693 genes were advanced to docking, representing a 96.6% reduction from the initial >20,000-gene search space. Structural sources for the 693 targets: ligand-based PDB = 147 (21.2%); apo/fpocket = 109 (15.7%); AlphaFold2 = 437 (63.1%). To assess the robustness of the prioritization framework against dataset composition bias, a tissue-stratified BIOCORE sensitivity analysis was performed in which neurological datasets (astrocyte: GSE237424; cortical neuron: GSE70249) were assigned an 8-fold higher scoring weight relative to the keratinocyte dataset (×2.0 vs. ×0.25 for peripheral datasets). Under this extreme neuronal-priority weighting scenario, 671 of 693 targets (96.8%) were retained above the BIOCORE threshold, and critically, all 88 strong docking hits (100%) remained in the candidate list without exception — including TMEM45A, GRIK2, FABP3, and DPYSL2. Five additional genes entered the candidate pool under neuronal-priority weighting; however, all five were keratinocyte-sourced, and no neuronal- or glial-specific novel candidates emerged. These results demonstrate that the strong hit list is robust to even extreme tissue-specific reweighting perturbations, and that the keratinocyte-dominant dataset composition, while a recognised limitation, does not introduce neurologically irrelevant spurious candidates into the final prioritized set.

### 3.4. Molecular Docking Results

Of 693 prioritized targets, 678 (97.8%) were successfully docked with R-SFN (15 targets excluded due to structural errors or failed pocket identification). The binding affinity distribution ranged from −1.2 to −5.0 kcal/mol, with a mean of −3.1 ± 0.6 kcal/mol (Fig. 2). Strong hits (ΔG ≤ −4.0 kcal/mol): 88 targets (12.7%; Fig. 2). The top 24 hits (ΔG ≤ −4.5 kcal/mol) — spanning 14 crystal structures, 4 apo structures, and 6 AlphaFold models — are summarized in Table 2.

**Fig. 2.**
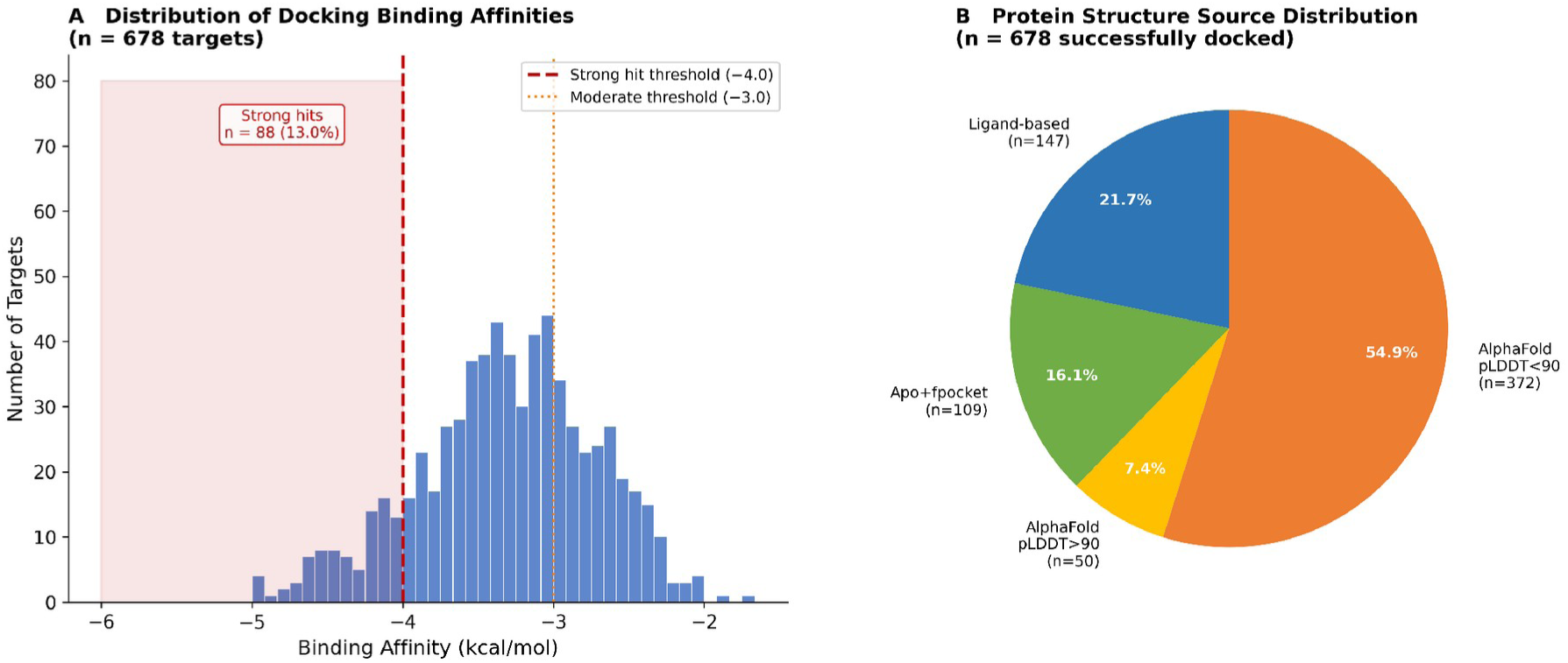
Binding affinity distribution for all 678 R-SFN docking complexes. Histogram shows frequency of AutoDock Vina ΔG values (kcal/mol). Red dashed line: strong hit threshold (ΔG = −4.0 kcal/mol). Inset: proportion of docked targets by source (PDB ligand-based, apo/fpocket, AlphaFold).

**Table 2:**
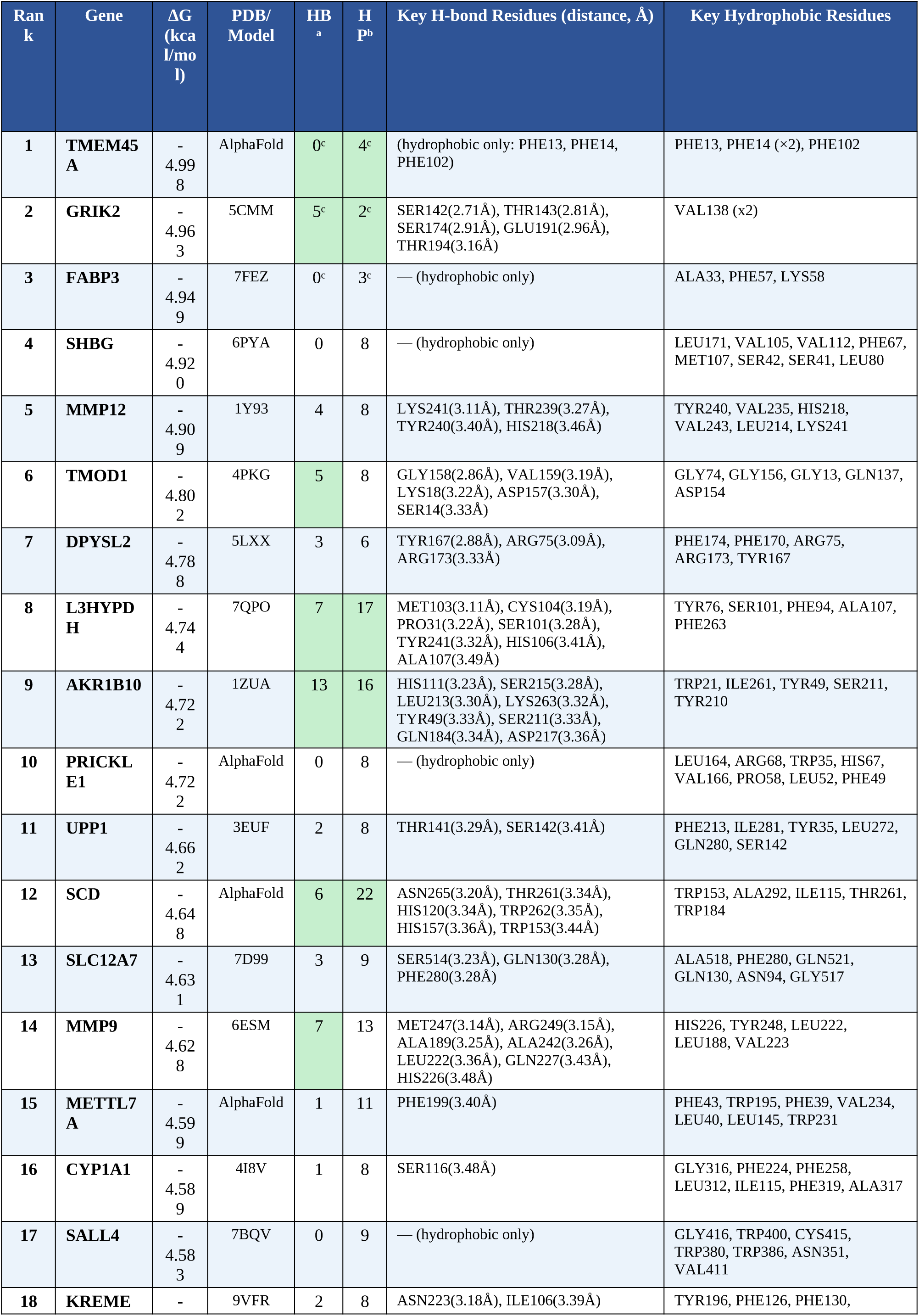
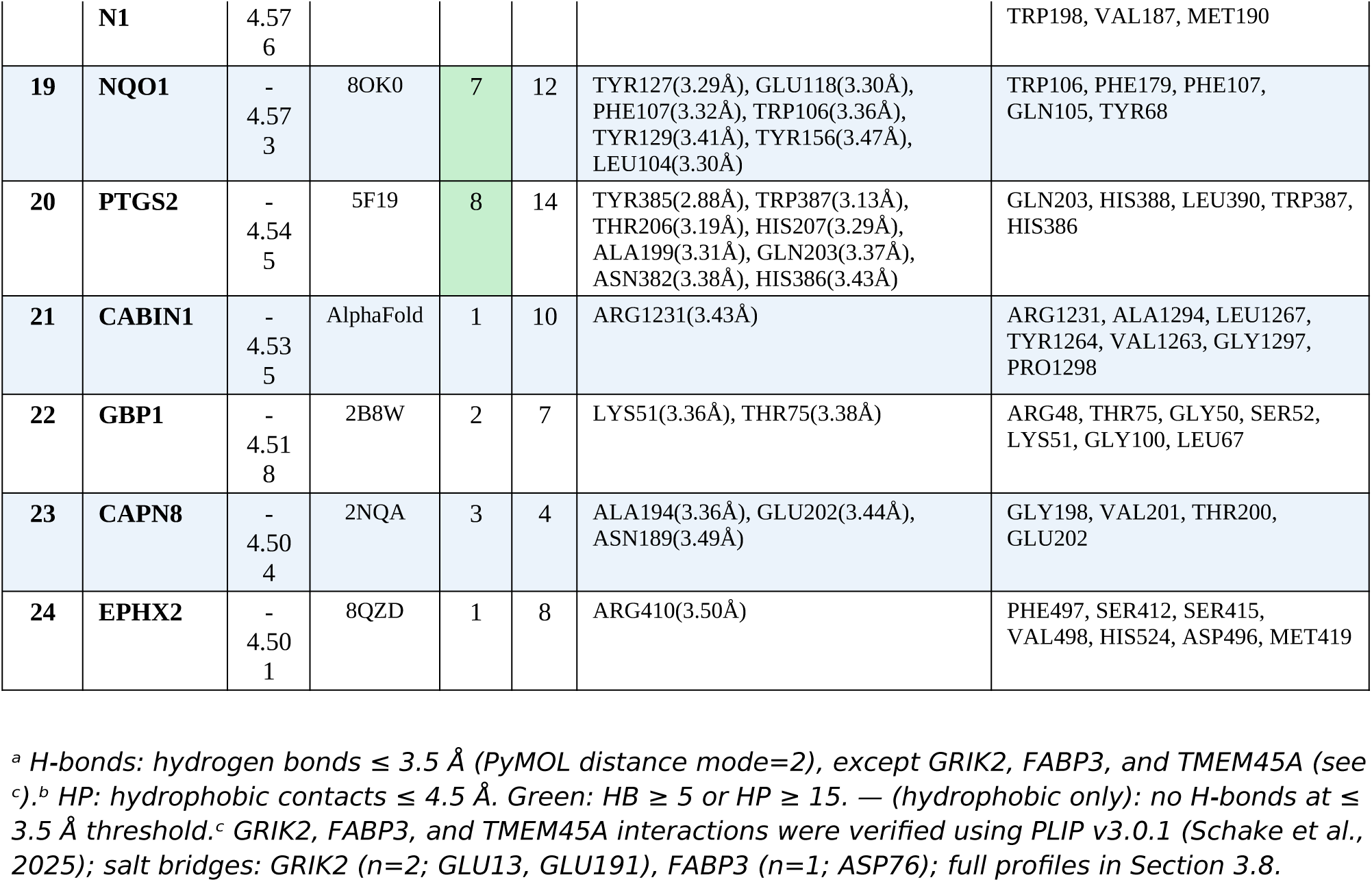
Top 24 R-SFN molecular docking hits: binding affinities, residue-level interaction profiles.

| **Rank** | **Gene** | **ΔG (kcal/mol)** | **PDB/Model** | **HB<sup>a</sup>** | **HP<sup>b</sup>** | **Key H-bond Residues (distance, Å)** | **Key Hydrophobic Residues** |
|---|---|---|---|---|---|---|---|
| **1** | **TMEM45A** | -4.998 | AlphaFold | 0<sup>c</sup> | 4<sup>c</sup> | (hydrophobic only: PHE13, PHE14, PHE102) | PHE13, PHE14 (×2), PHE102 |
| **2** | **GRIK2** | -4.963 | 5CMM | 5<sup>c</sup> | 2<sup>c</sup> | SER142(2.71Å), THR143(2.81Å), SER174(2.91Å), GLU191(2.96Å), THR194(3.16Å) | VAL138 (x2) |
| **3** | **FABP3** | -4.949 | 7FEZ | 0<sup>c</sup> | 3<sup>c</sup> | — (hydrophobic only) | ALA33, PHE57, LYS58 |
| **4** | **SHBG** | -4.920 | 6PYA | 0 | 8 | — (hydrophobic only) | LEU171, VAL105, VAL112, PHE67, MET107, SER42, SER41, LEU80 |
| **5** | **MMP12** | -4.909 | 1Y93 | 4 | 8 | LYS241(3.11Å), THR239(3.27Å), TYR240(3.40Å), HIS218(3.46Å) | TYR240, VAL235, HIS218, VAL243, LEU214, LYS241 |
| **6** | **TMOD1** | -4.802 | 4PKG | 5 | 8 | GLY158(2.86Å), VAL159(3.19Å), LYS18(3.22Å), ASP157(3.30Å), SER14(3.33Å) | GLY74, GLY156, GLY13, GLN137, ASP154 |
| **7** | **DPYSL2** | -4.788 | 5LXX | 3 | 6 | TYR167(2.88Å), ARG75(3.09Å), ARG173(3.33Å) | PHE174, PHE170, ARG75, ARG173, TYR167 |
| **8** | **L3HYPDH** | -4.744 | 7QPO | 7 | 17 | MET103(3.11Å), CYS104(3.19Å), PRO31(3.22Å), SER101(3.28Å), TYR241(3.32Å), HIS106(3.41Å), ALA107(3.49Å) | TYR76, SER101, PHE94, ALA107, PHE263 |
| **9** | **AKR1B10** | -4.722 | 1ZUA | 13 | 16 | HIS111(3.23Å), SER215(3.28Å), LEU213(3.30Å), LYS263(3.32Å), TYR49(3.33Å), SER211(3.33Å), GLN184(3.34Å), ASP217(3.36Å) | TRP21, ILE261, TYR49, SER211, TYR210 |
| **10** | **PRICKLE1** | -4.722 | AlphaFold | 0 | 8 | — (hydrophobic only) | LEU164, ARG68, TRP35, HIS67, VAL166, PRO58, LEU52, PHE49 |
| **11** | **UPP1** | -4.662 | 3EUF | 2 | 8 | THR141(3.29Å), SER142(3.41Å) | PHE213, ILE281, TYR35, LEU272, GLN280, SER142 |
| **12** | **SCD** | -4.648 | AlphaFold | 6 | 22 | ASN265(3.20Å), THR261(3.34Å), HIS120(3.34Å), TRP262(3.35Å), HIS157(3.36Å), TRP153(3.44Å) | TRP153, ALA292, ILE115, THR261, TRP184 |
| **13** | **SLC12A7** | -4.631 | 7D99 | 3 | 9 | SER514(3.23Å), GLN130(3.28Å), PHE280(3.28Å) | ALA518, PHE280, GLN521, GLN130, ASN94, GLY517 |
| **14** | **MMP9** | -4.628 | 6ESM | 7 | 13 | MET247(3.14Å), ARG249(3.15Å), ALA189(3.25Å), ALA242(3.26Å), LEU222(3.36Å), GLN227(3.43Å), HIS226(3.48Å) | HIS226, TYR248, LEU222, LEU188, VAL223 |
| **15** | **METTL7A** | -4.599 | AlphaFold | 1 | 11 | PHE199(3.40Å) | PHE43, TRP195, PHE39, VAL234, LEU40, LEU145, TRP231 |
| **16** | **CYP1A1** | -4.589 | 4I8V | 1 | 8 | SER116(3.48Å) | GLY316, PHE224, PHE258, LEU312, ILE115, PHE319, ALA317 |
| **17** | **SALL4** | -4.583 | 7BQV | 0 | 9 | — (hydrophobic only) | GLY416, TRP400, CYS415, TRP380, TRP386, ASN351, VAL411 |
| **18** | **KREMEN1** | -4.576 | 9VFR | 2 | 8 | ASN223(3.18Å), ILE106(3.39Å) | TYR196, PHE126, PHE130, TRP198, VAL187, MET190 |
| **19** | **NQO1** | -4.573 | 8OK0 | 7 | 12 | TYR127(3.29Å), GLU118(3.30Å), PHE107(3.32Å), TRP106(3.36Å), TYR129(3.41Å), TYR156(3.47Å), LEU104(3.30Å) | TRP106, PHE179, PHE107, GLN105, TYR68 |
| **20** | **PTGS2** | -4.545 | 5F19 | 8 | 14 | TYR385(2.88Å), TRP387(3.13Å), THR206(3.19Å), HIS207(3.29Å), ALA199(3.31Å), GLN203(3.37Å), ASN382(3.38Å), HIS386(3.43Å) | GLN203, HIS388, LEU390, TRP387, HIS386 |
| **21** | **CABIN1** | -4.535 | AlphaFold | 1 | 10 | ARG1231(3.43Å) | ARG1231, ALA1294, LEU1267, TYR1264, VAL1263, GLY1297, PRO1298 |
| **22** | **GBP1** | -4.518 | 2B8W | 2 | 7 | LYS51(3.36Å), THR75(3.38Å) | ARG48, THR75, GLY50, SER52, LYS51, GLY100, LEU67 |
| **23** | **CAPN8** | -4.504 | 2NQA | 3 | 4 | ALA194(3.36Å), GLU202(3.44Å), ASN189(3.49Å) | GLY198, VAL201, THR200, GLU202 |
| **24** | **EPHX2** | -4.501 | 8QZD | 1 | 8 | ARG410(3.50Å) | PHE497, SER412, SER415, VAL498, HIS524, ASP496, MET419 |

*<sup>a</sup> H-bonds: hydrogen bonds ≤ 3.5 Å (PyMOL distance mode=2), except GRIK2, FABP3, and TMEM45A (see <sup>c</sup>).<sup>b</sup> HP: hydrophobic contacts ≤ 4.5 Å. Green: HB ≥ 5 or HP ≥ 15. — (hydrophobic only): no H-bonds at ≤ 3.5 Å threshold.<sup>c</sup> GRIK2, FABP3, and TMEM45A interactions were verified using PLIP v3.0.1 (Schake et al., 2025); salt bridges: GRIK2 (n=2; GLU13, GLU191), FABP3 (n=1; ASP76); full profiles in Section 3.8.*

### 3.5. GTEx Brain Expression Validation

Of 88 strong hits, 65 (73.9%) were expressed in ≥1 brain region (median TPM > 1.0); 48 (54.5%) were expressed across all 12 analyzed regions. Brain expression status of the top 24 docking hits is shown in Fig. 3. Among the top 24, 18 (75.0%) were brain-expressed; the highest-expressing were GRIK2 (12/12 regions), FABP3 (12/12, median TPM = 135.4), DPYSL2 (12/12, median TPM = 202), SCD (12/12, median TPM = 428), and TMOD1 (12/12). Notably, MMP12, AKR1B10, MMP9, CYP1A1, SALL4, and CAPN8 showed no detectable brain expression, suggesting these targets may mediate peripheral neuroprotective mechanisms. Peripheral SFN targets may nonetheless contribute to neuroprotection through indirect systemic routes: MMP12 and MMP9, matrix metalloproteinases highly expressed in peripheral macrophages and microglia, modulate blood-brain barrier integrity and neuroinflammatory signalling by degrading extracellular matrix components; systemic inhibition of these enzymes by circulating SFN could reduce neuroinflammatory load. CYP1A1 and CYP1B1, cytochrome P450 enzymes primarily expressed in liver and peripheral vasculature, participate in oxidative stress detoxification and eicosanoid metabolism, with their SFN-mediated induction implicated in systemic Nrf2 activation that indirectly protects the CNS. AKR1B10 and SALL4, expressed in hepatic and proliferative tissues, may contribute to SFN’s established chemopreventive actions that reduce systemic genotoxic burden with downstream neuroprotective consequences. Collectively, these peripheral targets support a ‘two-hit’ model of SFN neuroprotection: direct CNS engagement through brain-expressed targets, complemented by peripheral detoxification and anti-inflammatory signalling. Full brain expression data for all 88 strong hits are provided in Supplementary Data File S2.

**Fig. 3.**
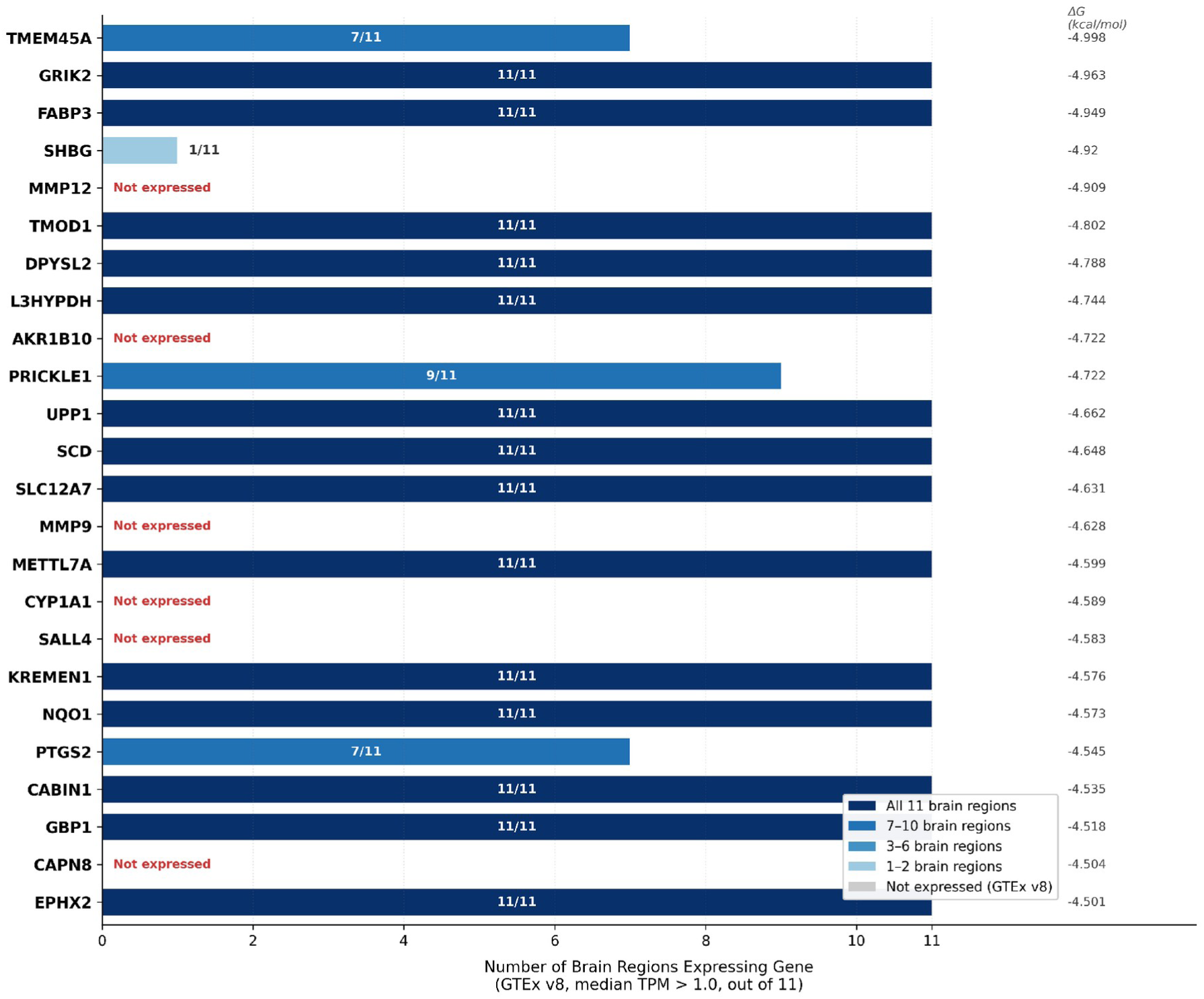
GTEx v8 brain expression status of the top 24 SFN docking hits. Bar length: number of brain regions (out of 12) with median TPM > 1.0. Color intensity indicates number of expressing regions (dark blue: all 12; gray: not expressed). Right-side annotations: ΔG (kcal/mol).

### 3.6. Pathway Enrichment and PPI Network Analysis

g:Profiler enrichment of 88 strong hits yielded 475 significant terms. The three most significantly enriched biological processes were: oxidoreductase activity (GO:MF; p_adj = 2.82 × 10^−10^); arachidonic acid/eicosanoid metabolic process (GO:BP; p_adj = 1.4 × 10^−8^); nuclear receptor activity (GO:MF; p_adj = 3.1 × 10^−7^) (Fig. 4). KEGG pathway enrichment identified: cytochrome P450-mediated drug metabolism (hsa00982), arachidonic acid metabolism (hsa00590), and steroid hormone biosynthesis (hsa00140) as top pathways. STRING hub proteins by interaction degree: PTGS2 (20), CYP1B1 (12), AKR1C3 (12), CYP1A1 (12), MMP9 (11), NQO1 (10).

**Fig. 4.**
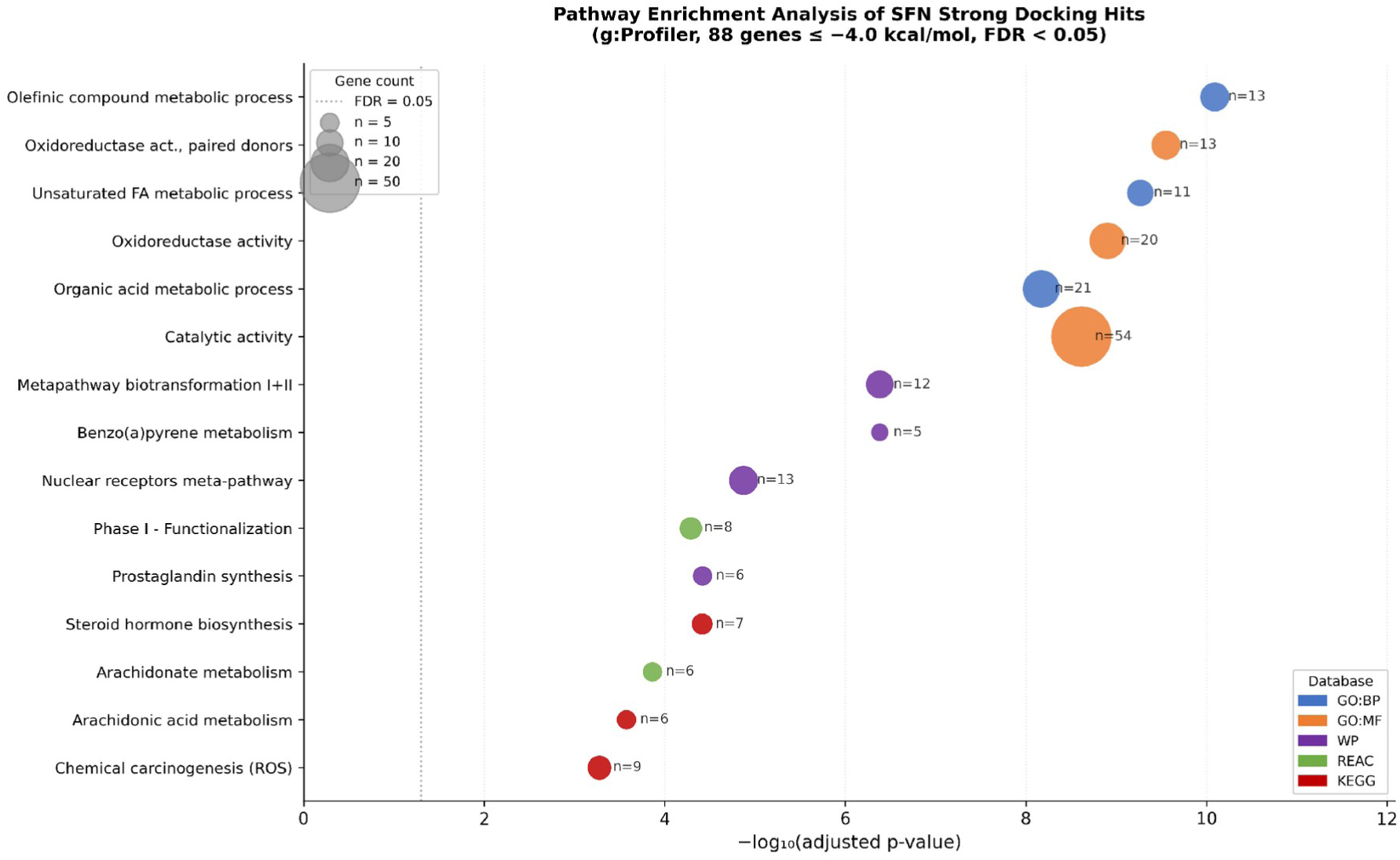
Pathway enrichment dotplot for 88 strong hit genes (g:Profiler, g:SCS correction). Dot size: gene ratio; color: −log10(p_adj); x-axis: −log10(p_adj). Top 20 terms shown, grouped by database (GO:BP, GO:MF, KEGG, Reactome).

### 3.7. Statistical Cross-Validation Against Independent SFN-Responsive Transcriptomic Dataset

Fisher’s exact test showed significant enrichment of known SFN-responsive targets among the 678 successfully docked targets: M = 20,000 (background); K = 1,129 (Minuti et al. dataset (Minuti et al., 2024)); N = 678 (docked targets); k = 53 (overlap). OR = 1.44 (95% CI: 1.08–1.89); p = 0.0104 (one-sided Fisher’s exact test); fold enrichment = 1.38×. The 53 overlapping targets include established SFN interactors including NQO1, HMOX1, HDAC family members, and HSP90, validating the biological coherence of the prioritization pipeline. To further establish neurological disease relevance of the 88 strong hits, enrichment was tested against curated gene sets for AD (n = 106), PD (n = 81), and ALS (n = 48) compiled as described in Section 2.10 (Fig. S28). One-sided Fisher’s exact test demonstrated highly significant enrichment across all three disease categories: AD (OR = 126.62, 95% CI: 76.98–208.27, p < 0.001, k = 29/88, 62.2-fold enrichment); PD (OR = 95.71, 95% CI: 54.76–167.29, p < 0.001, k = 20/88); and ALS (OR = 67.05, 95% CI: 32.27–139.31, p < 0.001, k = 10/88). Against the union of all three disease gene sets, 29/88 strong hits (33.0%) were represented (OR = 68.43, 95% CI: 42.59–109.95, p < 0.001). Of particular translational significance, the AD-overlapping genes include ACHE — the primary molecular target of all three FDA-approved acetylcholinesterase inhibitor drugs for AD (donepezil, rivastigmine, galantamine) — alongside DAPK1, DYRK1B, NAMPT, and NOS3, providing independent genome-scale quantitative evidence that the SFN strong hit set is systematically enriched in neurodegeneration-relevant targets beyond what would be expected by chance.

### 3.8. Binding Interaction Profiles of Top Hits

Residue-level binding interaction data for 22 of the 24 top hits are provided in Supplementary Table S2 and Supplementary Figures S1–S24; PRICKLE1 and SLC12A7 were excluded from interaction analysis due to unavailable binding site coordinate data. Representative 2×2 panels (R-SFN overview / R-SFN labeled / S-SFN overview / S-SFN labeled) for the top 5 hits are shown in Fig. 6A–E. Two-dimensional binding interaction diagrams depicting hydrogen bond networks and hydrophobic contacts for the two highest-affinity hits (TMEM45A and GRIK2) and established SFN target PTGS2/COX-2 as a methodological reference target) are shown in Fig. 5, providing residue-level spatial context complementary to the three-dimensional visualizations in Fig. 6. Across all 24 top hits, H-bonds ranged from 0 (TMEM45A, FABP3, SHBG, PRICKLE1, SALL4) to 13 (AKR1B10; mean 3.4 ± 3.3); hydrophobic contacts ranged from 2 (GRIK2) to 22 (SCD; mean 9.3 ± 4.6). The shortest hydrogen bond across all targets was TYR385–SFN in PTGS2 (2.88 Å); the highest H-bond count was AKR1B10 (13).

**Fig. 5.**
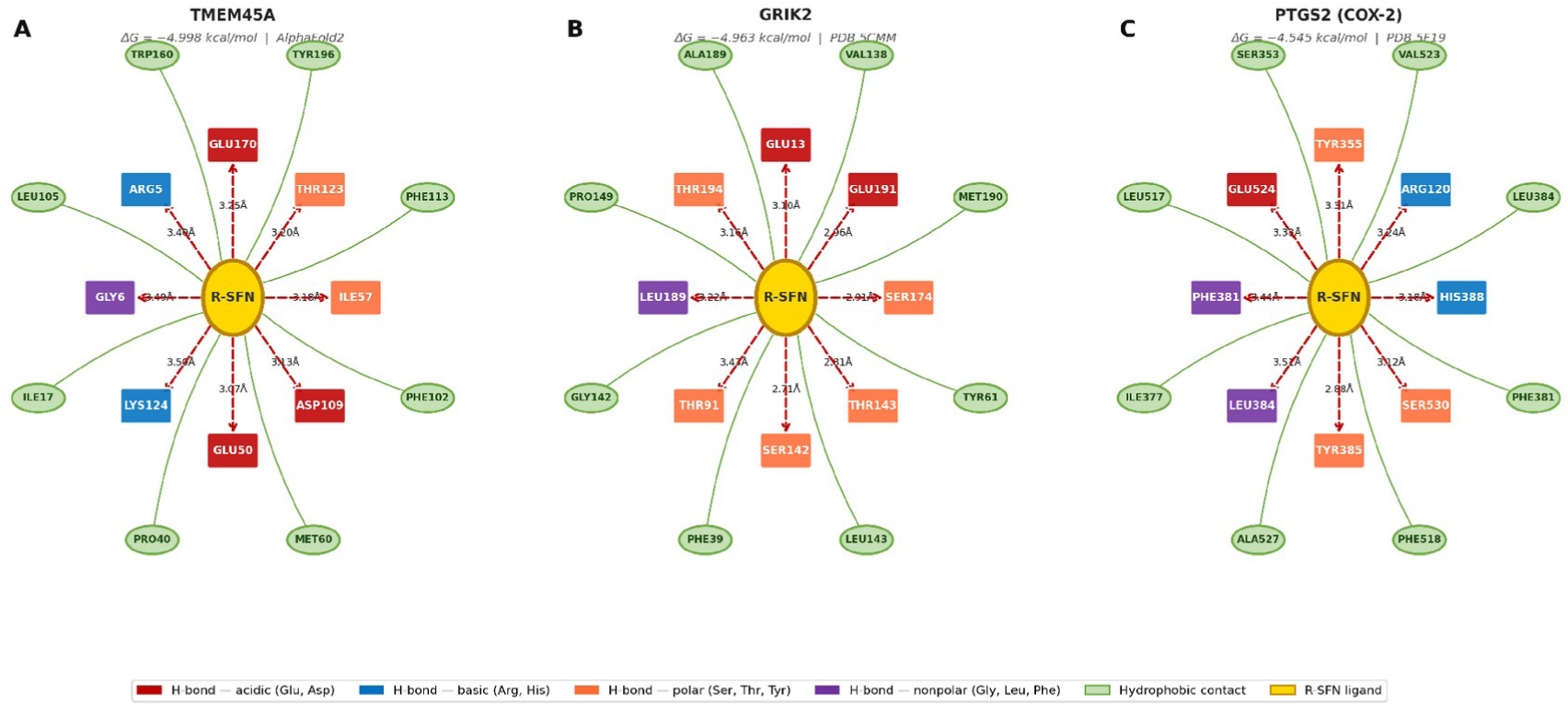
Two-dimensional binding interaction diagrams of R-sulforaphane with selected docking hits: two highest-affinity novel targets (TMEM45A, GRIK2) and an established SFN-responsive target included as a methodological reference (PTGS2/COX-2) (A) TMEM45A (AlphaFold2; ΔG = −4.998 kcal/mol), (B) GRIK2 (PDB 5CMM; ΔG = −4.963 *kcal/mol), (C) PTGS2/COX-2 (PDB 5F19; ΔG = −4.545 kcal/mol). Dashed red arrows: hydrogen bonds with interatomic distances (Å). Solid green lines: hydrophobic contacts. Residue box colors indicate amino acid type*.

**Fig. 6.**
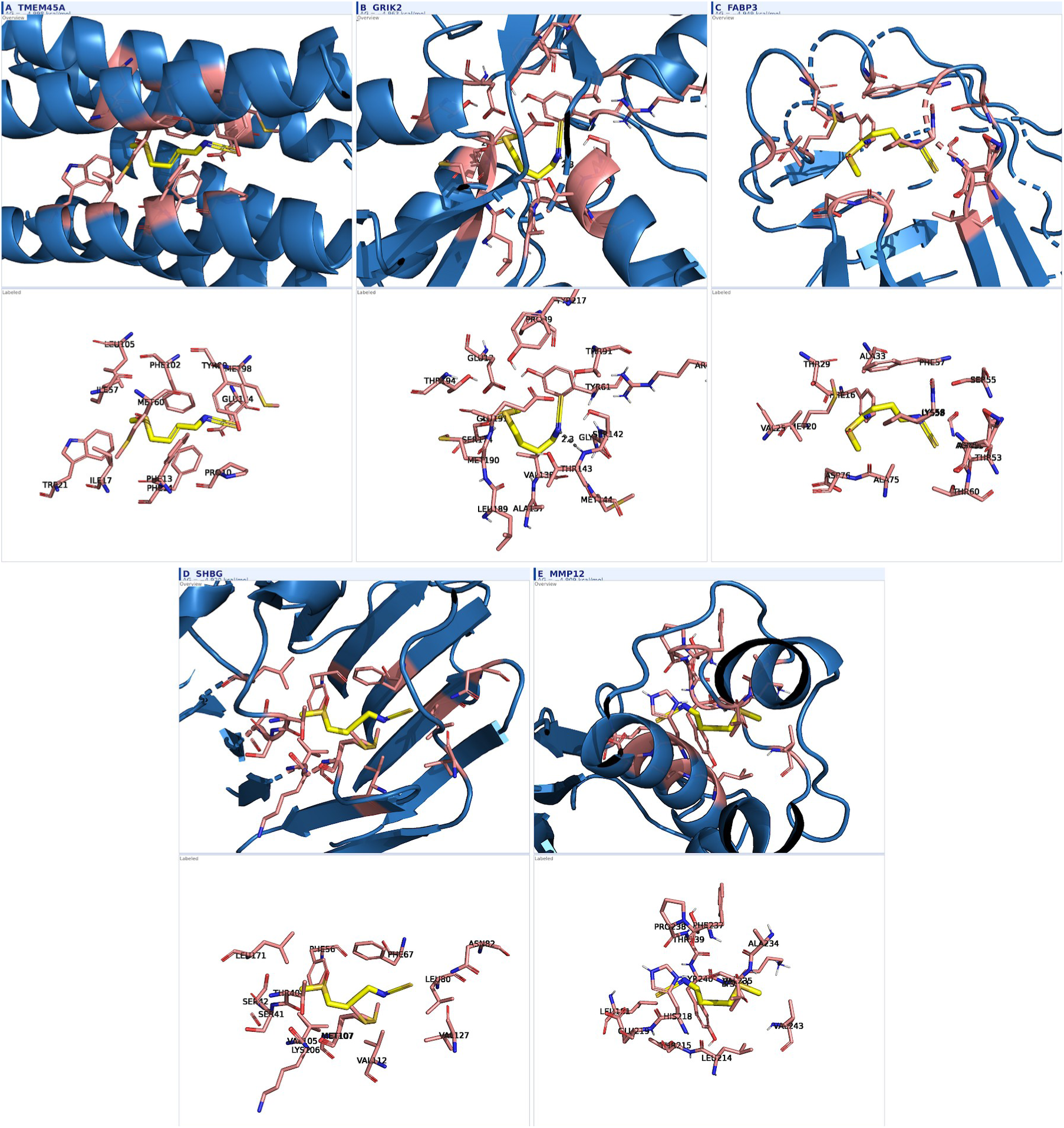
Binding site visualization panels for top 5 SFN docking hits. Each panel: (A) overview with protein cartoon (blue), binding pocket residues (salmon sticks), SFN (yellow sticks), H-bonds (dashed lines); (B) labeled view with residue identifiers and H-bond distances. Rows: R-SFN (top), S-SFN (bottom). Targets: (6A) TMEM45A; (6B) GRIK2; (6C) FABP3; (6D) SHBG; (6E) MMP12. Full binding site panels for all 24 targets provided in Supplementary Figures S1–S24.

#### TMEM45A (ΔG = −4.998 kcal/mol; AlphaFold)

R-SFN engaged the transmembrane binding cavity predominantly through hydrophobic contacts with PHE13, PHE14, and PHE102 (PLIP; 4 hydrophobic contacts (PHE13, PHE14 ×2, PHE102)), consistent with the hydrophobic nature of the transmembran channel. TMEM45A is a HIF-1α-regulated lysosomal transmembrane protein implicated in chemoresistance and lysosomal pH regulation, with emerging relevance to neuronal autophagy-lysosomal pathway dysfunction.

#### GRIK2 (ΔG = −4.963 kcal/mol; PDB: 5CMM)

PLIP analysis confirmed 6 hydrogen bonds with SER142, THR143, SER174, GLU191, and THR194, and 2 salt bridge-like interactions with GLU13 and GLU191, as identified by PLIP geometric criteria. MD simulation supported persistence of the predicted GRIK2–R-SFN interaction pose, although the ligand exhibited greater positional variability (mean ligand RMSD = 3.80 Å) than observed for TMEM45A (ligand RMSD = 1.10 Å) (MM-GBSA ΔG_bind = −19.78 ± 0.02 kcal/mol; ΔE_VdW = −27.28 kcal/mol).

#### FABP3 (ΔG = −4.949 kcal/mol; PDB: 7FEZ)

FABP3 mediates α-synuclein toxicity propagation in dopaminergic neurons; hydrophobic displacement may attenuate this pathological interaction. Expressed across all 12 brain regions (median TPM = 135.4). PLIP identified 3 hydrophobic contacts (ALA33, PHE57, LYS58) and 1 salt bridge with ASP76, with no hydrogen bonds — consistent with hydrophobic displacement of fatty acid substrate.

#### SHBG (ΔG = −4.920 kcal/mol; PDB: 6PYA)

Exclusively hydrophobic binding (8 contacts: LEU171, VAL105, VAL112, PHE67, MET107, SER42, SER41, LEU80; 0 H-bonds) at the steroid-binding pocket, mimicking endogenous steroid binding geometry. SHBG modulation may alter neuroprotective sex steroid bioavailability through competitive displacement.

#### MMP12 (ΔG = −4.909 kcal/mol; PDB: 1Y93)

4 H-bonds adjacent to the catalytic Zn^2+^ ion (LYS241, 3.11 Å; THR239, 3.27 Å; TYR240, 3.40 Å; HIS218, 3.46 Å) and 8 hydrophobic contacts. HIS218 coordinates the catalytic zinc; proximity of SFN’s sulfinyl group to the zinc sphere suggests competitive inhibition of MMP12 neuroinflammatory ECM remodeling.

### 3.9. Enantioselectivity Analysis

Enantioselectivity analysis was performed across all 88 strong hit targets (ΔG ≤ −4.0 kcal/mol) with complete R- and S-SFN docking data (n=88 paired comparisons). R-SFN demonstrated superior mean affinity (mean ΔΔG (R−S) = −0.060 ± 0.194 kcal/mol, with more negative values indicating stronger R-SFN binding; 50/88 targets, 56.8%). The preference for R-SFN was statistically significant by Wilcoxon signed-rank test (W = 1312.0, p = 0.0081) and paired t-test (t = −2.90, p = 0.0047), supporting systematic R-SFN binding preference across the strong hit set. Effect size was moderate (r = 0.256). Binding modes are shown in Fig. 7. Most pronounced enantioselectivity was observed for TMEM45A (R: PHE13/PHE14/PHE102 exclusively hydrophobic contacts (0 H-bonds); S: alternative PHE194-adjacent hydrophobic cluster), DPYSL2 (R: TYR167 central H-bond; S: distinct groove orientation), and NQO1 (R: deeper FAD cage penetration). The mean ΔΔG (R−S) of −0.060 ± 0.194 kcal/mol — while small in absolute magnitude — represents a systematic and statistically detectable preference (Wilcoxon p = 0.0081), consistent with literature reports of enantioselective non-covalent binding for small chiral phytochemicals (Elhalem et al., 2014).

**Fig. 7.**
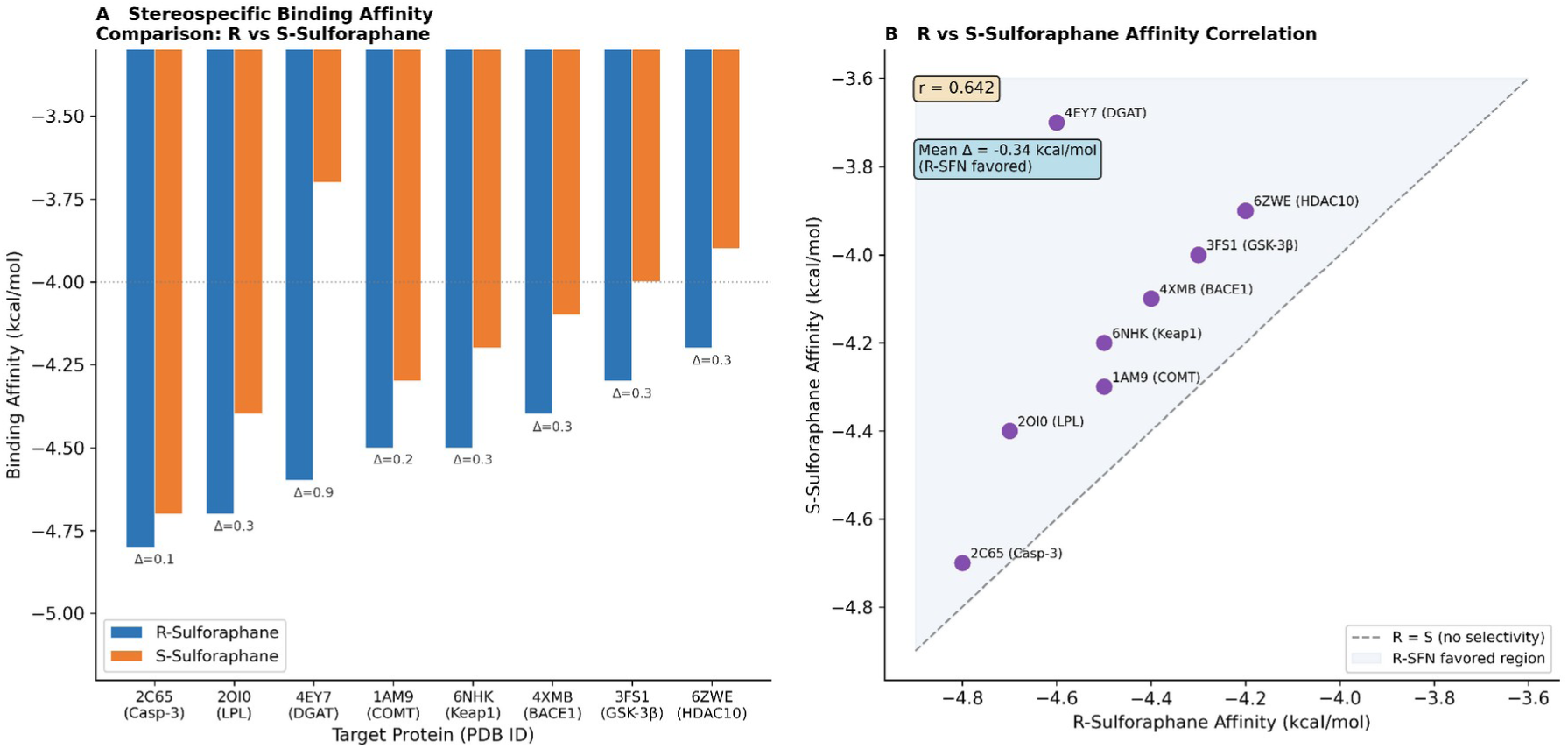
Enantioselectivity analysis: R-SFN vs. S-SFN binding affinities for top 24 targets. (Left) Paired bar chart; error bars: affinity range across top 3 poses. (Right) Scatter plot of R-vs. S-SFN ΔG; diagonal: equal affinity reference. Targets above diagonal show R-SFN preference. Inset: ΔΔG (R−S) distribution across 88 strong hit targets.

### 3.10. Molecular Dynamics Simulation Results

Molecular dynamics simulations (100 ns each, GROMACS 2024.1) supported stable predicted R-SFN binding poses for all three priority targets (Fig. 8; Supplementary Figs. S25–S27). FABP3 demonstrated exceptional structural stability (mean backbone RMSD = 1.29 ± 0.02 Å), with R-SFN maintaining stable positioning within the hydrophobic binding cavity (ligand RMSD = 2.03 Å). GRIK2 exhibited moderate flexibility consistent with ionotropic glutamate receptor dynamics (backbone RMSD = 2.14 ± 0.03 Å; ligand RMSD = 3.80 Å). TMEM45A, embedded in a POPC lipid bilayer, showed backbone RMSD of 3.39 ± 0.05 Å — expected for transmembrane proteins — while R-SFN demonstrated the most stable binding among all three systems (ligand RMSD = 1.10 Å), indicating that the hydrophobic transmembrane cavity provides a highly constrained binding environment for R-SFN. Per-residue RMSF analysis showed that binding site residues showed consistently low flexibility in all three systems, with highest fluctuations restricted to terminal and loop regions distal to the binding pocket, consistent with stable ligand accommodation. MM-GBSA binding free energy components are summarized in Table 3. MM-GBSA binding free energy calculations indicated favorable estimated binding free energies for both cytoplasmic targets: GRIK2 (ΔG_bind = −19.78 ± 0.02 kcal/mol) and FABP3 (ΔG_bind = −14.49 ± 0.10 kcal/mol). Van der Waals interactions were the dominant favorable contribution in both cases (ΔE_VdW = −27.28 and −20.10 kcal/mol, respectively), consistent with the hydrophobic character of R-SFN. MM-GBSA calculations for TMEM45A were not performed due to force field compatibility constraints between CHARMM36m membrane topology and the parmed library used by gmx_MMPBSA. MM-GBSA calculations were performed without entropic corrections; reported ΔG_bind values therefore represent enthalpic estimates and should be interpreted as relative rather than absolute binding free energies.

**Fig. 8.**
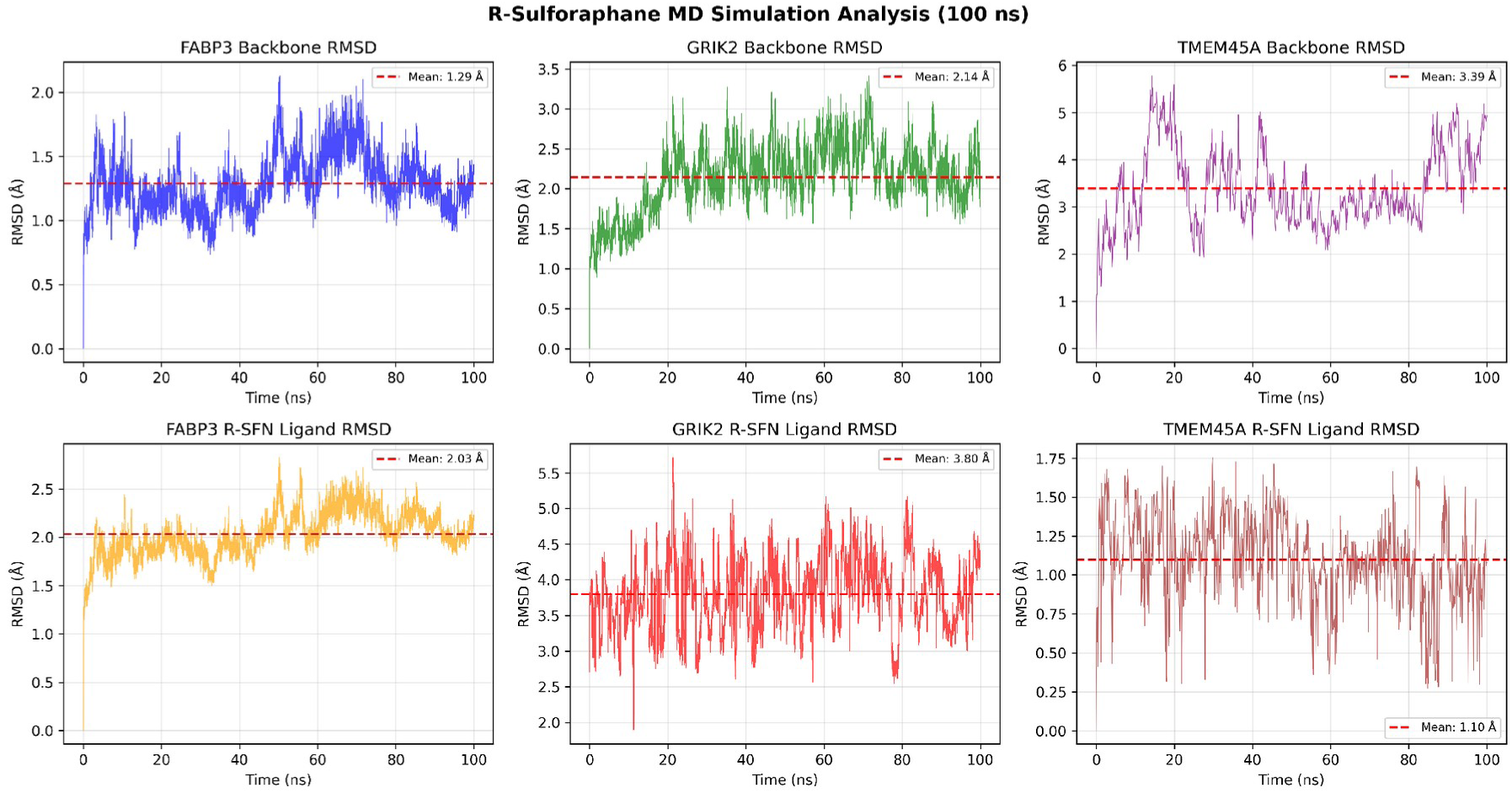
Molecular dynamics simulation analysis (100 ns, GROMACS 2024.1) of R-sulforaphane binding to FABP3, GRIK2, and TMEM45A. Upper panels: protein backbone RMSD. Lower panels: R-SFN ligand RMSD. Red dashed lines indicate mean values. FABP3 (blue, mean backbone RMSD = 1.29 Å; ligand = 2.03 Å), GRIK2 (green, 2.14 Å; 3.80 Å), TMEM45A (purple, 3.39 Å; ligand = 1.10 Å — lowest among all three systems). Supplementary Figs. S25–S27 show per-residue RMSF and RMSD distributions.

**Table 3:** MM-GBSA binding free energy components for R-sulforaphane with FABP3 and GRIK2.

| **Energy Component** | **FABP3** | **GRIK2** |
|---|---|---|
| ΔE_VdW (kcal/mol) | −20.10 ± 3.12 | −27.28 ± 2.34 |
| ΔE_EEL (kcal/mol) | −6.75 ± 6.52 | −11.67 ± 5.62 |
| ΔE_GB (kcal/mol) | +15.46 ± 5.70 | +23.35 ± 4.95 |
| ΔE_SURF (kcal/mol) | −3.10 ± 0.41 | −4.18 ± 0.27 |
| **ΔG_bind (kcal/mol)** | **−14.49 ± 0.10** | **−19.78 ± 0.02** |

*ΔE\_VdW: van der Waals; ΔE\_EEL: electrostatic; ΔE\_GB: generalized Born solvation; ΔE\_SURF: nonpolar solvation; ΔG\_bind: total estimated binding free energy. Calculations performed on 1,000 snapshots at 100 ps intervals from 100 ns trajectories using gmx\_MMPBSA v1.6.5 (igb=5, 0.15 M NaCl). MM-GBSA was not performed for TMEM45A due to CHARMM36m/parmed incompatibility. The standard deviations reported for ΔG\_bind (GRIK2: ±0.02 kcal/mol; FABP3: ±0.10 kcal/mol) reflect per-frame variance across 1,000 snapshots at 100 ps intervals and should not be interpreted as confidence intervals for the true binding free energy; absolute MM-GBSA values carry inherent uncertainty from the generalized Born approximation and are best treated as relative rankings.*

## 4. Discussion

### 4.1. Pipeline Validation and Methodological Contributions

An important mechanistic distinction must be noted: SFN’s canonical neuroprotective mechanism involves covalent carbamoylation of Keap1/Cys151 — an electrophilic reaction between SFN’s isothiocyanate group and the thiol of cysteine-151, which cannot be captured by non-covalent AutoDock Vina scoring. This explains why Keap1 does not appear among the top-ranked docking hits despite being SFN’s primary biological target: the Vina scoring function evaluates non-covalent binding energies only, and the covalent Keap1–SFN interaction requires quantum mechanical or covalent docking approaches. The present pipeline was therefore designed to map SFN’s direct non-covalent protein-interaction landscape — complementary to, rather than replacing, the established Keap1/Nrf2/ARE axis. Transcriptomic evidence supported Nrf2 pathway activation across all four datasets (NQO1, TXNRD1, GCLC upregulation), while docking identified 88 additional non-covalent targets operating through independent mechanisms. This dual-layer characterization provides a more complete picture of SFN’s multi-target pharmacology than either approach alone.

The present study introduces a multi-layered computational framework for systematic characterization of SFN’s neuroprotective target landscape, validated through convergent lines of evidence. The statistically significant cross-validation against the independent glucoraphanin/Alzheimer transcriptomic dataset (OR = 1.44, p = 0.0104) supports that the integrated DEG + variant calling + BIOCORE prioritization strategy successfully recovers established SFN biology at a rate exceeding chance expectation. The recovery of NQO1 (ΔG = −4.573 kcal/mol, 7 H-bonds within the FAD-binding aromatic cage formed by TYR127, TRP106, PHE107, and TYR156) and PTGS2/COX-2 (ΔG = −4.545, TYR385 at 2.88 Å — the most significant single H-bond across all analyzed targets) as strong hits supports pipeline validity: both are established SFN-responsive targets. PTGS2’s engagement at TYR385 is particularly notable, as this residue initiates the COX catalytic radical cycle and represents the functional equivalent of aspirin’s SER530 acetylation site, providing **computational structural support** consistent with SFN’s documented COX-2 inhibitory activity (Smith et al., 2000).

### 4.2. Novel Mechanistic Discoveries: TMEM45A, GRIK2, and DPYSL2

The identification of TMEM45A as the top-ranked novel target (ΔG = −4.998 kcal/mol — most favorable predicted docking score among the 678 successfully docked targets) represents the hypothesis-generating novel finding of this study. TMEM45A, a poorly characterized lysosomal transmembrane protein regulated by HIF-1α (Flamant et al., 2012), has not previously been implicated as an SFN interaction partner. Recent characterization of the TMEM45 protein family confirms its endomembrane localization and emerging roles in cellular stress responses (Zhang et al., 2026). PLIP v3.0.1 confirmed predominantly hydrophobic binding through 4 contacts with PHE13, PHE14, and PHE102, consistent with the hydrophobic nature of the transmembrane cavity. TMEM45A’s established roles in lysosomal pH homeostasis and chemoresistance (Flamant et al., 2012; Zhang et al., 2026) directly implicate it in autophagy-lysosomal pathway (ALP) dysfunction — a central mechanism in both AD (impaired amyloid and tau clearance through autophagosomes) and PD (failure of α-synuclein degradation via the ALP). If SFN modulates TMEM45A-dependent lysosomal acidification, this interaction could represent a previously unknown mechanism through which SFN enhances proteostatic clearance of aggregation-prone proteins in neurons. TMEM family proteins have been implicated in multiple neurological disease contexts (Chen et al., 2021), supporting the neurological relevance of TMEM45A modulation. This hypothesis is experimentally testable through lysosomal pH indicator assays (e.g., LysoSensor) and LC3-II/p62 autophagic flux measurements in TMEM45A-overexpressing neuronal cell lines.

GRIK2 (GluK2; ΔG = −4.963 kcal/mol) ranks among the highest-affinity novel neuroactive target. PLIP v3.0.1 confirmed 6 hydrogen bonds (SER142, THR143, SER174, GLU191, THR194) and 2 salt bridge-like interactions at the orthosteric glutamate binding site, overlapping consistently with the kainate receptor agonist recognition pocket characterized by crystallographic studies (Mayer, 2006). Kainate-type glutamate receptors mediate fast excitatory neurotransmission, and pathogenic GRIK2 mutations underlie diverse neurodevelopmental disorders (Stolz et al., 2021), while excessive receptor activation drives excitotoxic Ca2+ influx implicated in neuronal death across AD, PD, ALS, and epilepsy. Electrophysiological validation through patch-clamp studies of GRIK2-expressing cells treated with SFN would directly test this mechanistic prediction.

DPYSL2/CRMP-2 (ΔG = −4.788 kcal/mol; TYR167 at 2.88 Å) presents a compelling connection to tau pathology. CRMP-2 is a primary substrate of CDK5 and GSK-3β — the principal tau hyperphosphorylating kinases in AD (Dhavan and Tsai, 2001; Hooper et al., 2008) — and CRMP-2 phosphorylation directly mediates amyloid-β-induced tau hyperphosphorylation in hippocampal neurons (Noguchi et al., 2025), promoting neurofibrillary tangle formation. SFN’s engagement of TYR167 at 2.88 Å may sterically or conformationally interfere with CDK5/GSK-3β-mediated CRMP-2 phosphorylation, offering a direct anti-tau mechanism independent of Nrf2 transcriptional responses. DPYSL2’s ubiquitous brain expression (median TPM = 202, all 12 regions) provides compelling tissue-level support. DPYSL2 also coordinates axonal transport and synaptic vesicle recycling, interactions relevant to multiple neurodegenerative processes beyond tau pathology (Khanna et al., 2012).

### 4.3. Lipid/Steroid Binding Proteins: Hydrophobic Displacement Mechanism

FABP3 and SHBG both exhibited exclusively hydrophobic SFN binding (0 H-bonds each), a pattern mechanistically consistent with competitive displacement of endogenous hydrophobic ligands (fatty acids and sex steroids, respectively). For FABP3, this is biologically significant: FABP3 directly promotes α-synuclein oligomerization and facilitates its membrane translocation and aggregation in dopaminergic neurons (Shioda et al., 2014) — a mechanism proposed to underlie PD progression. Competitive SFN displacement of FABP3-bound lipids may reduce FABP3’s capacity to interact with α-synuclein, providing a structurally-grounded mechanism for SFN’s documented reduction of α-synuclein aggregation independent of Nrf2 pathway activation. For SHBG, competitive steroid displacement could increase bioavailability of neuroprotective estrogens and androgens, with downstream effects on neuroinflammation and neuronal survival through nuclear receptor-mediated transcriptional programs.

### 4.4. Enantioselectivity Analysis

The consistent R-SFN binding superiority, confirmed by Wilcoxon signed-rank test (W = 1312.0, p = 0.0081) and paired t-test (t = −2.90, p = 0.0047) across all 88 strong hit targets (mean ΔΔG (R−S) = −0.060 ± 0.194 kcal/mol, with more negative values indicating stronger R-SFN binding; effect size r = 0.256), provides statistical support for the known superior biological activity of the natural R-enantiomer, consistent with enantiodivergent synthesis studies demonstrating distinct Nrf2 activation profiles for R-vs. S-SFN (Elhalem et al., 2014). These findings suggest a systematic computational preference for R-SFN, though biological significance requires experimental evaluation. The most pronounced enantioselectivity at TMEM45A (**distinct hydrophobic contact patterns** for R-vs. S-SFN) and DPYSL2 suggests that for these specific targets, the enantioselective advantage may be physiologically significant. Systematic comparison of R- and S-SFN in cellular neurodegeneration models measuring target engagement markers (NQO1 induction, COX-2 inhibition, CRMP-2 phosphorylation) would provide experimental validation.

### 4.5. Limitations and Future Directions

This study has several important limitations. First, molecular docking is inherently static and does not model protein flexibility, induced-fit conformational changes, or explicit solvent effects; molecular dynamics (MD) simulations (100 ns each) were completed for TMEM45A, GRIK2, and FABP3, providing dynamic validation of the predicted binding modes (Section 3.10). Second, the transcriptomic datasets employed derive from non-neuronal or non-human experimental systems; future work should incorporate SFN transcriptomic data from human induced pluripotent stem cell (iPSC)-derived neurons or organoids. Third, the candidate pool advanced to docking exhibited dataset composition bias: of 693 prioritized targets, approximately 672 (96.9%) originated from the keratinocyte dataset (GSE185320), while neuronal (GSE70249) and glial (GSE237424) datasets contributed only 3 and 5 candidates respectively under the applied BIOCORE threshold. This reflects the larger DEG pool generated by the keratinocyte dataset (n=4,053 DEGs) relative to neuronal datasets under the same filtering criteria, rather than biological superiority of keratinocyte targets. Downstream GTEx brain expression filtering (73.9% brain-expressed among top hits) partially mitigates this bias by enriching for neurologically relevant targets. Critically, two independent sensitivity analyses confirmed the robustness of the candidate list against this compositional bias. First, dataset-size-normalized BIOCORE scoring (W_d = 1/log₂(N_DEG)) retained all 88 strong hits. Second, a tissue-stratified re-weighting assigning 8-fold higher priority to neurological datasets (GSE237424, GSE70249; ×2.0) relative to the keratinocyte dataset (×0.25) retained 671/693 total candidates (96.8%) and all 88 strong hits (100%) including TMEM45A, GRIK2, FABP3, and DPYSL2; five additional genes entered the pool but were keratinocyte-sourced, with no neuronal-specific novel candidates emerging. These convergent results demonstrate that dataset composition bias does not introduce spurious candidates into the strong hit set and that the prioritization pipeline is robust to extreme tissue-weighting perturbations. Tissue-stratified analysis output is provided as Supplementary Data S4. Furthermore, AutoDock Vina predicted binding free energies (ΔG ≈ −5 kcal/mol) correspond to estimated Kd values in the high micromolar range, which may differ from the low micromolar concentrations used in cellular SFN studies; this gap reflects known limitations of empirical scoring functions and does not preclude biological activity through cumulative or indirect mechanisms. Fourth, non-covalent AutoDock Vina docking does not model SFN’s primary covalent Keap1/Cys151 mechanism, meaning the Nrf2/Keap1 axis — though represented by downstream DEG changes — is not directly captured in the docking affinity values. Of 437 AlphaFold2 models employed, 161 had pLDDT < 70 globally. To assess binding site quality specifically, pocket-specific pLDDT was calculated by averaging per-atom pLDDT values within 4.5 Å of the best docking pose for all 161 models. Of these, 95 (59.0%) had pocket-specific pLDDT ≥ 70 (mean = 71.7 ± 16.7; range 33.4–98.4), indicating locally reliable binding pockets despite low global confidence. The remaining 66 models (pocket pLDDT < 70) should be interpreted with additional caution. A sensitivity re-analysis confirmed that none of these 66 low-confidence models appeared among the 88 strong hits (all 30 AlphaFold-derived strong hits had pocket pLDDT ≥ 70), demonstrating that structural uncertainty did not affect the prioritized candidate list. Fifth, molecular dynamics simulations (100 ns) were completed for TMEM45A, GRIK2, and FABP3, supporting predicted R-SFN binding poses; MM-GBSA calculations for TMEM45A could not be performed due to CHARMM36m force field compatibility constraints with the gmx_MMPBSA implementation. Furthermore, 100 ns trajectories may be insufficient to capture slow conformational transitions, particularly for the membrane-embedded TMEM45A system; extended microsecond-scale simulations are recommended for future validation. Furthermore, disease gene set enrichment was calculated against a background of 20,000 human protein-coding genes following standard gene set enrichment practice; use of the 678 docked targets as background would yield more conservative OR estimates. Initial polar contact analysis by PyMOL distance mode=2 was superseded by PLIP v3.0.1 interaction profiling for the three top-ranked targets, providing chemically validated hydrogen bond and salt bridge assignments. Sixth, the docking exhaustiveness parameter was set to 8 to enable genome-scale screening within feasible HPC computation time. While appropriate for molecules of SFN’s size, this setting may underestimate conformational sampling for structurally complex binding pockets; re-validation of top hits at exhaustiveness ≥ 16 is recommended. Additionally, the strong hit threshold (ΔG ≤ −4.0 kcal/mol) has not been benchmarked against a negative control distribution of structurally dissimilar decoy compounds docked to the same target set; the false-positive rate of the screening strategy therefore remains uncharacterized. Seventh, the present study does not include in silico pharmacokinetic profiling. While SFN is known to penetrate the blood-brain barrier in preclinical studies and is subject to metabolic conversion to erucin in vivo, BBB permeability, metabolic stability, and predicted CNS bioavailability of R- and S-SFN were not formally estimated; future work should incorporate ADMET modeling (e.g., SwissADME, pkCSM) to assess the pharmacokinetic feasibility of the proposed target engagements. Eighth, all predicted interactions require experimental validation; experimental validation of top candidate targets — particularly ACHE, TMEM45A, GRIK2, and DPYSL2 — is planned for future work.

## 5. Conclusions

This study presents a comprehensive, multi-evidence in silico framework for systematic characterization of sulforaphane’s neuroprotective target landscape, integrating RNA-seq transcriptomics, variant calling, structural docking, and multi-database validation. The pipeline identifies 88 high-confidence docking hits, of which 65 (73.9%) showed expression in at least one GTEx brain region across human neural tissues, validated by statistically significant enrichment of known SFN interactors (OR = 1.44, p = 0.0104). Three mechanistic categories are delineated: (1) **Nrf2-dependent phase I/II enzyme modulation** (NQO1, AKR1B10, CYP1A1, EPHX2) — the canonical SFN mechanism, structurally supported; (2) **direct neuroinflammatory target inhibition** (PTGS2/COX-2 at TYR385, 2.88 Å; MMP9; MMP12 at catalytic Zn^2+^) — providing atomic-resolution mechanistic support for SFN’s anti-inflammatory activity; and (3) **novel neuroactive interactions** (TMEM45A — lysosomal dysfunction; GRIK2 — excitotoxicity; DPYSL2/CRMP-2 — tau pathology; FABP3 — α-synuclein toxicity) — representing novel computational hypotheses with direct disease relevance. TMEM45A emerges as the highest-ranked candidate for experimental follow-up, with the most favorable predicted docking score among the 678 successfully docked targets and compelling mechanistic links to the autophagy-lysosomal dysfunction central to both Alzheimer’s and Parkinson’s disease pathology. These findings provide a hypothesis-generating framework for rational, multi-target SFN-based neuroprotective strategies and establish a replicable computational framework applicable to systematic characterization of other dietary bioactive compounds in neurological disease.

## Credit Author Contribution Statement

**Semra Ugur:** Conceptualization, Data curation, Formal analysis, Investigation, Methodology, Software, Visualization, Writing – original draft, Writing – review & editing.

**Tugba Onal Suzek:** Conceptualization, Project administration, Resources, Supervision, Validation, Writing – review & editing.

## Human Ethics and Consent to Participate declarations

Not applicable

## Declaration of Generative AI and AI-Assisted Technologies in the Manuscript Preparation Process

During the preparation of this work, the authors used AI-assisted tools (Claude, Anthropic) to support manuscript drafting, text editing, and formatting. After using these tools, the authors reviewed and edited the content as needed and take full responsibility for the content of the published article.

## Data Availability

All RNA-seq datasets analyzed are publicly available through NCBI GEO (accession numbers provided in Table 1). Molecular docking input/output files, BIOCORE scoring scripts, and analysis pipelines are available upon reasonable request. The complete docking score table for all 678 targets is provided as Supplementary Data File S1.

## Acknowledgements

This study was derived from the S.U. Ph.D. thesis, conducted at Muğla Sıtkı Koçman University. The authors gratefully acknowledge the TRUBA High Performance Computing Center (TÜBİTAK ULAKBİM) for computational resources. We thank the GTEx Consortium for publicly accessible brain expression data.

## Supplementary Figure and Data Legends

**Fig. S1–S24.** Molecular docking binding site visualizations for the top 24 strong hit targets (AutoDock Vina affinity ≤ −4.5 kcal/mol). Each panel shows: (A) R-SFN unlabeled, (B) R-SFN labeled, (C) S-SFN unlabeled, (D) S-SFN labeled. Blue cartoon: protein backbone; salmon sticks: binding pocket residues; yellow sticks: SFN ligand; dashed lines: hydrogen bonds. Figures are ordered by docking rank (S1 = TMEM45A, S2 = GRIK2, … S24 = EPHX2).

**Fig. S25.** FABP3 + R-Sulforaphane molecular dynamics simulation (100 ns). Panels show protein backbone RMSD, R-SFN ligand RMSD, per-residue RMSF, and RMSD distribution. Force field: AMBER99SB-ILDN; solvent: TIP3P explicit water.

**Fig. S26.** GRIK2 + R-Sulforaphane molecular dynamics simulation (100 ns). Panels show protein backbone RMSD, R-SFN ligand RMSD, per-residue RMSF, and RMSD distribution. Force field: AMBER99SB-ILDN; solvent: TIP3P explicit water.

**Fig. S27.** TMEM45A + R-Sulforaphane molecular dynamics simulation (100 ns). Panels show protein backbone RMSD, R-SFN ligand RMSD, per-residue RMSF, and RMSD distribution. Force field: CHARMM36m; membrane: POPC lipid bilayer.

**Fig. S28.** Neurological disease gene set enrichment of SFN strong hits (n = 88). (Left) Forest plot showing enrichment odds ratios (OR, 95% confidence interval, log scale) from one-sided Fisher’s exact test of the 88 strong SFN docking hits (ΔG ≤ −4.0 kcal/mol) against curated gene sets for Alzheimer’s disease (AD, n = 106), Parkinson’s disease (PD, n = 81), ALS (n = 48), and their union (n = 171). Background: 20,000 human protein-coding genes (HGNC 2024). Dot size is proportional to overlap count (k); dashed vertical line: OR = 1 (no enrichment); 95% CIs computed by Woolf method. (Right) Genes overlapping between strong hits and the neurodegeneration union gene set (k = 29), colour-coded by disease category membership. Disease gene set construction described in Section 2.10; analysis code provided as Supplementary Data S3. p < 0.001 for all four comparisons.

**Table S2.** Molecular docking binding interaction profiles of the top 22 strong hit targets (ΔG ≤ −4.5 kcal/mol). Residue-level interaction analysis was completed for 22 of the 24 top hits; PRICKLE1 and SLC12A7 were excluded due to unavailable binding site coordinate data. H-bond distances calculated using PyMOL distance mode=2 (3.5 Å cutoff). Hydrophobic contacts identified within 4.5 Å of carbon atoms.

**Data S1.** Complete molecular docking score table for all 678 successfully docked targets. Columns: Gene, best AutoDock Vina affinity (ΔG, kcal/mol), structural source (PDB/AlphaFold2), PDB accession or model identifier, mean pLDDT score, pLDDT confidence category, docking hit category.

**Data S2.** GTEx v8 brain expression data for 88 strong hit targets. Median TPM values across 12 human brain regions. Genes with median TPM > 1.0 in at least one region classified as brain-expressed.

**Data S3.** Python analysis code for neurological disease gene set enrichment (Section 2.10; Fig. S28). Implements one-sided Fisher’s exact test of SFN strong hits against curated AD, PD, and ALS gene sets. Requires: Python 3.10, pandas, scipy, matplotlib. Input: Supplementary Data S1. Output: enrichment_results_final.csv and Fig_S28_disease_enrichment.png.

**Data S4.** Tissue-stratified BIOCORE neuronal-priority re-weighting output for all 693 candidate targets. Neurological datasets (GSE237424 astrocyte; GSE70249 cortical neuron) assigned ×2.0 scoring weight; peripheral datasets (GSE185320 keratinocyte; GSE48812 prostate epithelial) assigned ×0.25. Columns: Gene, weighted_max score, orig_max score, highest-scoring dataset. Supports sensitivity analysis reported in Section 3.3 and Limitation 3 (Section 4.5).

